# Sex-Specific Ethylene Responses Drive Floral Sexual Plasticity in Cannabis

**DOI:** 10.1101/2025.06.03.657661

**Authors:** Adrian S. Monthony, Julien Roy, Maxime de Ronne, Olivia Carlson, Susan J. Murch, Davoud Torkamaneh

**Affiliations:** Département de phytologie, Université Laval, Québec City, Québec, Canada; Institut de Biologie Intégrative et des Systèmes (IBIS), Université Laval, Québec, Canada; Centre de recherche et d’innovation sur les végétaux (CRIV), Université Laval, Québec, Canada; Institut intelligence et données (IID), Université Laval, Québec, Canada; Department of Chemistry, University of British Columbia, Kelowna, Canada

**Keywords:** cannabis, sex determination, transcriptomics, multi-omics, sexual plasticity, ethylene, silver thiosulfate, ethephon

## Abstract

*Cannabis sativa* L. exhibits remarkable sexual plasticity: both XX and XY individuals can undergo complete phenotypic sex reversal in response to ethylene modulation. While this phenomenon is well documented, the molecular mechanisms remain underexplored. Here, we present the first multi-omic study of hormonally induced sex change in both XX and XY *Cannabis* plants, integrating transcriptomic profiling, ethylene pathway metabolite quantification, and whole-genome sequencing across three genetically distinct genotypes. Treatments with silver thiosulfate (STS) and ethephon induced >80% phenotypic conversion, but transcriptomic responses diverged sharply between chromosomal sexes. We profiled 47 ethylene-related genes (ERGs) and identified 14 high-confidence candidates—including *CsACS1*, *CsACO5*, *CsERF1*, and *CsMTN*—with sex-specific, time-dependent expression patterns that support a two-phase model of plasticity: early transcriptional reprogramming followed by stabilization of new floral identities. Several candidate ERGs were in non-recombining regions of the X chromosome or absent from the Y, while most showed low nucleotide diversity, suggesting functional constraint. These findings provide a high-resolution view of ethylene-responsive sex plasticity and demonstrate that convergent floral phenotypes arise from distinct regulatory programs in XX and XY plants. Our work advances the molecular understanding of sexual plasticity in dioecious species and identifies candidate genes for the development of sex-stable cultivars in *Cannabis* and other crops.

## Introduction

*Cannabis sativa* L. (cannabis) is diploid species (2n=20) with nine autosomes and one pair of XX/XY sex chromosomes (Lapierre et al. 2023a; Monthony et al. 2024). Cannabis produces indeterminate compound inflorescences that develop distinctly dimorphic flowers: female flowers (FFs) consist of an ovary, surrounded by two bracts, dotted with glandular trichomes and two pistils (Reed 1914; Spitzer-Rimon et al. 2019), while male flowers (MFs) are staminate, characterized by a segmented perianth and five hanging stamen (Reed 1914). The glandular trichomes found on FFs synthesize medicinally important cannabinoids, of which over 100 have been characterized and are of interest for their potential medicinal applications (Giese et al. 2015; Mudge et al. 2019). In contrast, MFs produce cannabinoids at significantly lower levels and are considered to have negligible value in medicinal or commercial cannabis applications (Welling et al. 2021). Most drug-type cannabis is dioecious, producing exclusively male (XY) or female (XX) flowers, though some monoecious plants, mostly industrial hemp, produce both on the same plant (Ferfuia et al. 2021). Unlike most species, where the presence or absence of the Y chromosome alone determines sex, cannabis exhibits remarkable sexual plasticity—a plant’s ability to alter its phenotypic sex in response to environmental or hormonal cues (Monthony et al. 2024). Under normal growth conditions the phenotypic sex of cannabis plants is concordant with sex chromosome karyotype. However, documented cases of hermaphroditism and monoecy have been reported to arise spontaneous and from human intervention (Kurtz et al. 2020; Punja and Holmes 2020). Although the cannabis genome has been sequenced and assembled (Grassa et al. 2021), and interest in sex determination is growing (Prentout et al. 2020; Adal et al. 2021; Shi et al. 2024b; Chen et al. 2025b; Orozco et al. 2025), the molecular pathways underlying sex expression remain poorly understood. Thus, understanding the genetic and hormonal regulation of flower development and sex determination in cannabis has significant implications for both basic plant biology and applied cultivation strategies.

Sexual plasticity, which has also been referred to as leaky, or labile sex expression (Käfer et al. 2022), has been described across diverse taxa of flowering monoecious and dioecious plants, including: *Carica papaya* (papaya; dioecious), *Spinacia oleracea* (spinach; dioecious), *Mercurialis annua* L. (monoecious) and *Amborella trichopoda* (dioecious; Komai and Masuda 2004; Sather et al. 2010; Lin et al. 2016; Anger et al. 2017; Cossard and Pannell 2021). However, the triggers that shift phenotypic sex or floral sex ratios differ widely among species and conditions, and include natural population variability (Anger et al. 2017), reproductive pressure (Ji et al. 2016; Cossard and Pannell 2021), abiotic or biotic stressors (Lin et al. 2016), and targeted molecular and plant growth regulator interventions (Thomas 2004; Zhang et al. 2017; García et al. 2020). Although the precise molecular mechanisms controlling sexual plasticity in cannabis remain poorly understood, anecdotal reports from underground cultivators and subsequent empirical validation by academic researchers strongly implicates ethylene biosynthesis and signaling (Green 2005; Lubell and Brand 2018; Moon et al. 2020; Jones and Monthony 2022). Known signaling inhibitors, such as silver thiosulfate (STS), reliably induce male flowers (IMFs) in XX cannabis plants, while treatment with the ethylene-releasing compound ethephon (commercially Ethrel®) consistently promotes female flower development (IFFs) in XY plants (Lubell and Brand 2018; Moon et al. 2020; Flajšman et al. 2021). Similar roles for ethylene have been demonstrated in the Cucurbitaceae family, where increases in endogenous ethylene or exogenous applications result in an increased ratio of female to male flowers, resulting in ethylene often being described as feminizing (Rajagopalan et al. 2004; Li et al. 2021). Research on Cucurbitaceae has resulted in well-characterized gene networks involving ethylene biosynthesis and signaling gene families, such as the ACC SYNTHASE (ACS), ACC OXIDASE (ACO), and ETHYLENE RESPONSE (ETR; Boualem et al. 2015; Zhang et al. 2017; Li et al. 2021). In *Cannabis*’s sister genus *Humulus* (hops), the dioecious species *H. lupulus* and *H. japonicus* also exhibit ethylene-mediated sexual plasticity, with similar responses to STS and ethephon being reported, but the molecular mechanisms underpinning this plasticity has not yet been elucidated (Akagi et al. 2025). Beyond these examples, the regulatory pathways underpinning ethylene-mediated sexual plasticity have not been widely studied, making the *C. sativa* a potentially interesting dioecious analogue to Cucurbitaceae. Recently, our research group presented a first integrative model of the ethylene precursor pathway, the Yang cycle (extensively review by Chen et al. 2025a), as well as ethylene biosynthesis and signaling pathways in cannabis (Monthony et al., 2024). This study generated an initial set of candidate ethylene-related genes (ERGs) hypothesized to govern sexual plasticity, supported by preliminary transcriptomic evidence and leveraging existing genetic datasets (Prentout et al. 2020; Adal et al. 2021). This model, as well as most studies of cannabis sex to date, have focused on the role of ERGs in IMF production, neglecting to consider whether XY feminization relies on reciprocal molecular and hormonal mechanisms.

Sexual plasticity is a complex, multi-layered process involving genetic, transcriptomic, hormonal, and metabolic networks. Previous transcriptomic studies, including those applying advanced inference methods like network ontology annotation (Orozco et al. 2025), have identified hormone-related gene expression changes associated with sexual plasticity and highlighted ethylene as an important regulator (Prentout et al. 2020; Adal et al. 2021). However, most research has primarily focused on sexual plasticity in XX plants and rarely explored reciprocal mechanisms in XY plants undergoing feminization, despite growing experimental evidence implicating ethylene as a central regulator of cannabis sex expression (Ram and Jaiswal 1970; Galoch 1978; Kurtz et al. 2020; Moon et al. 2020; Adal et al. 2021; Flajšman et al. 2021). Moreover, no previous studies have directly quantified ethylene-pathway metabolites in cannabis during sex transitions. Interpretation of these transcriptomic studies is further complicated by well-documented limitations of gene ontology (GO)-based methods, including incomplete annotations, inconsistent term specificity, and poor taxon representation—issues especially pronounced in non-model species like cannabis (Uygun et al. 2016; Gaudet and Dessimoz 2017; Fattel and Lawrence-Dill 2025). Another limitation of existing studies is the narrow focus on floral tissues during sex transition. This approach implicitly assumes that sex-determining signals arise only once flowers are visibly developing. However, in many plants species, key floral fate regulators—such as FLOWERING LOCUS T (FT)—are activated in leaves in response to photoperiod changes and then transported to the shoot apical meristem to trigger flowering (Tsuji 2017). Recently, evidence has emerged to support *C. sativa* following this conserved model, suggesting that *C. sativa* FT-like genes are responsive to photoperiod changes prior to the emergences of flowers (Dowling et al. 2024).

In this study, we apply multi-omic framework—integrating transcriptomic, metabolomic, and genomic data—to enable higher-resolution interrogation of temporally dynamic regulatory networks underlying sexual plasticity in cannabis. By analyzing both genotypic and pre-floral tissue variation, our study offers a more complete view of the ethylene-responsive signaling networks coordinating sexual plasticity in cannabis. Our findings reveal that this process is not governed by a single genetic switch but is instead a temporally dynamic process involving distinct sets of genes. We identify genes likely involved in the initiation of sex change shortly after treatment, as well as others that contribute to stabilizing the new phenotypic state. Together, these results support a dynamic, phase-dependent model of ethylene-regulated sexual plasticity in cannabis.

## Results

### Highly inducible sexual plasticity in XX and XY cannabis

The ability of cannabis to exhibit sexual plasticity was evaluated by quantifying the percentage of flowers in the inflorescence whose phenotypic sex is opposite to the plant’s chromosomal sex. A two-way fixed effects analysis of variance (ANOVA) was performed to assess the effects of treatment and genotype on the percentage of sex change, 28 days following the switch to a flowering photoperiod (12/12). Treatment had a significant effect on the extent of sex change (p < 0.001), whereas genotype (p = 0.07) and the interaction between genotype and treatment (p = 0.91) were not significant (Supplementary Table 9). Across all three genotypes, treated inflorescences exhibited average sex change rate > 80% (Table 1). In contrast, H_2_O treated control plants showed no evidence of sexual plasticity in any genotype (Table 1). The observed phenotypic changes in treated plants are visually presented in Figure 1 and Supplemental Figure 1.

**Figure 1.**
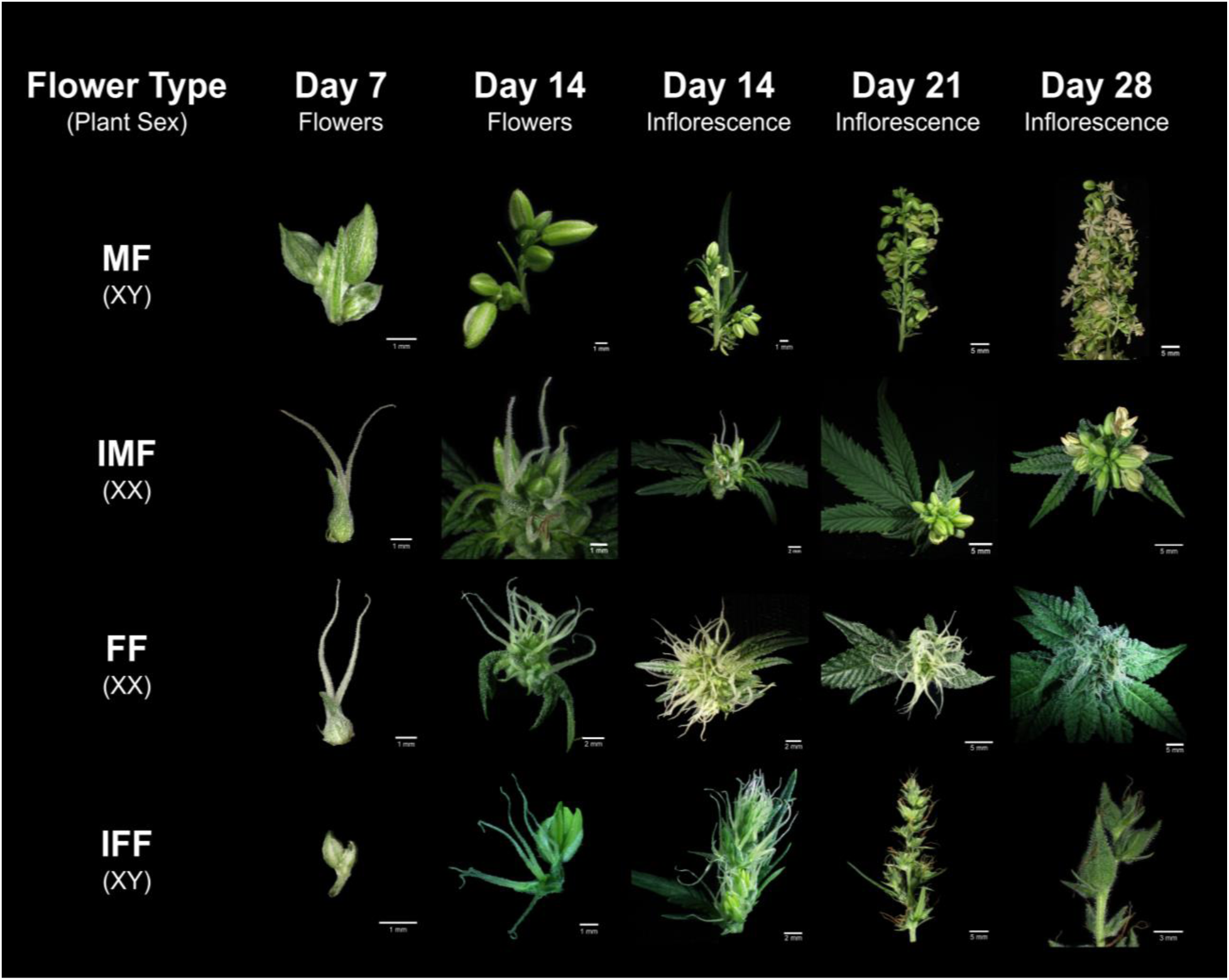
Representative timeline of floral development in MF, IMF, FF and IFF of *C. sativa* ‘La Rosca’ from 7 to 28 days after the photoperiod shift from long day to short day.

**Table 1.**
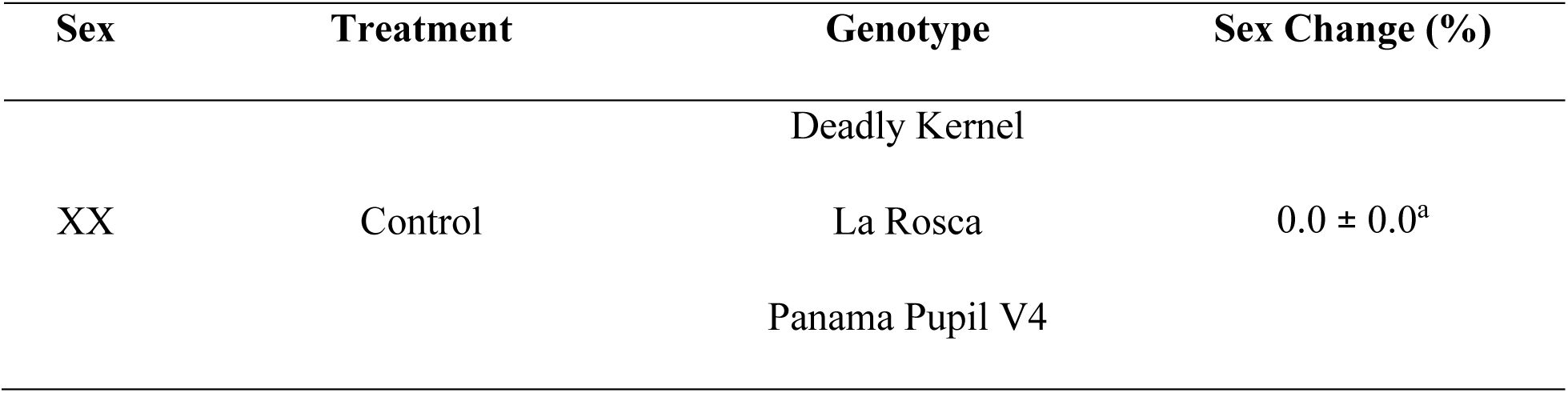
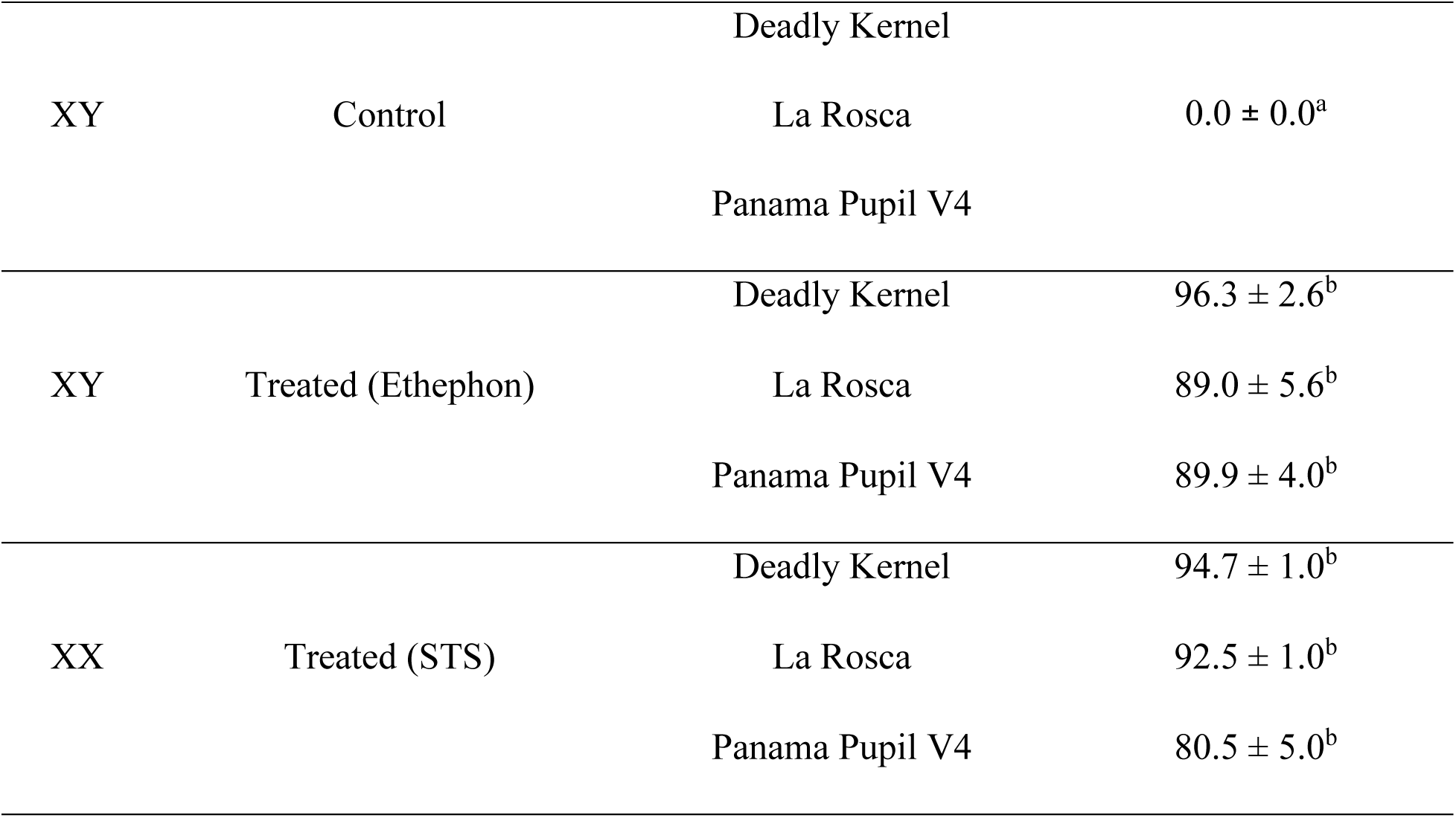
Percentage of florets in *C. sativa* inflorescences that changed sex in response to sex plasticity induction treatments. FF-female flowers; MF-male flowers; IFF-induced female flowers; IMF-induced male flowers. Uncertainty indicated by standard error of the mean (SEM) and differing letters indicate significant differences between treatments (p ≤ 0.05), as determined by a Tukey-Kramer post-hoc test.

Cannabinoid analysis of control female flowers obtained from the three XX genotypes confirmed THC-dominance (Chemotype I; Zheng 2022), with total THC content ranging from 16.98% to 25.40% w/w (Supplementary Table 10). Minor cannabinoids such as CBGA (0.28–1.20% w/w) and CBG (0.04–0.12% w/w) were also detected at varying levels (Supplementary Table 10).

### An updated ortholog identification reveals X-specific ERGs

An updated ortholog analysis identified six additional putative ERGs, expanding upon previous findings (Monthony et al. 2024). This analysis utilized newly available *C. sativa* genomes, including the new ‘Pink Pepper’ reference genome (NCBI 2023) and a fully phased X and Y-haplotype genomes from a male Otto II genotype (Carey et al. 2024b). In total, 47 ERGs were identified across the *C. sativa* genome (Supplementary Table 11), with the highest numbers found on chromosomes 1 and X (Figure 2A).

**Figure 2.**
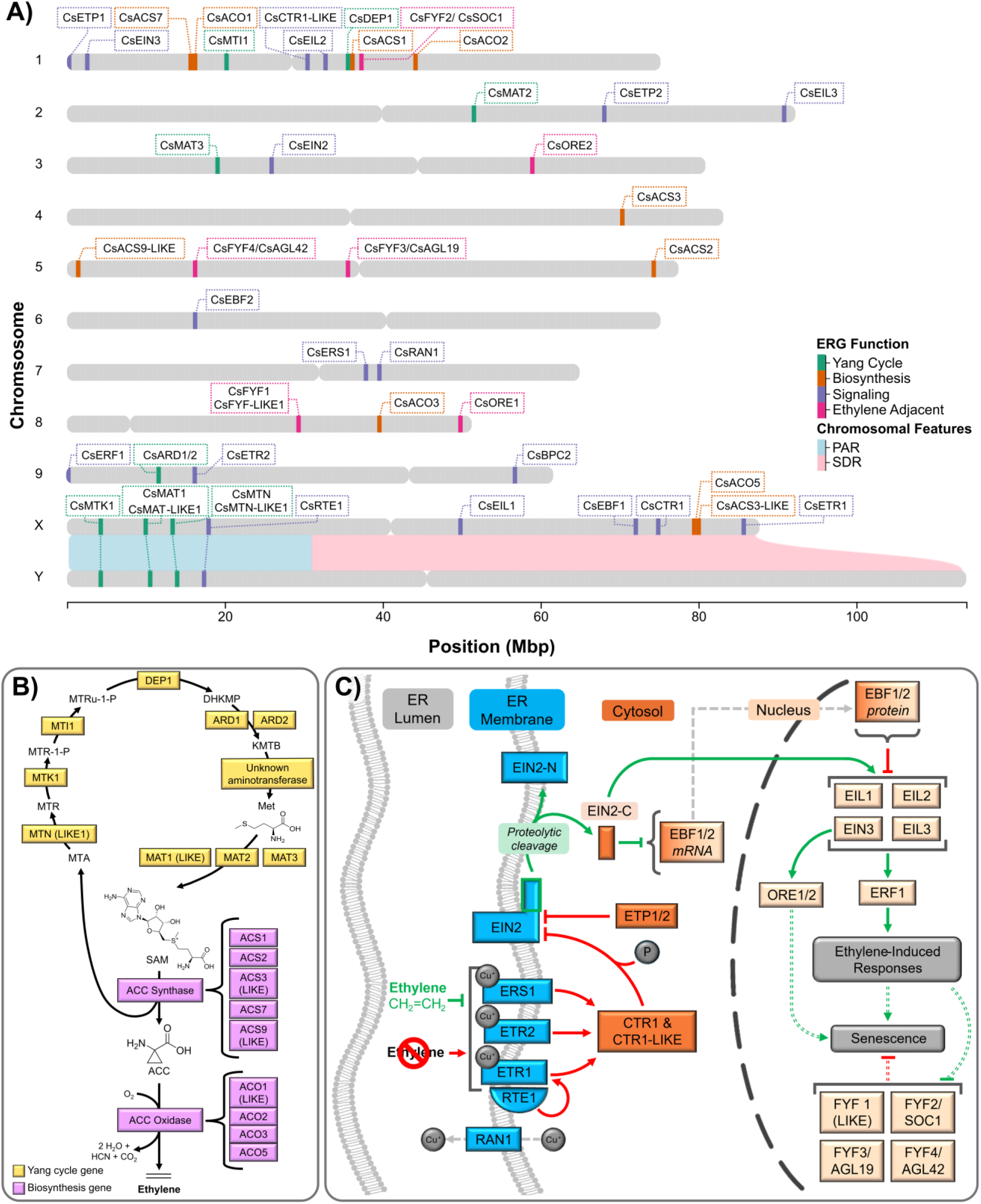
Updated genomic localization of ERGs and updated models of the ethylene pathway in *Cannabis sativa*. (A) Chromosomal positions of ethylene-related genes (ERGs) identified in *C. sativa* transcriptomes from Monthony et al. (2024) and expanded in the present study. Autosomal gene positions are based on the ‘Pink Pepper’ reference genome (NCBI, 2023); Pseudo-autosomal region (PAR) and non-recombining sex determination region (SDR) regions on the X and Y chromosomes follow Carey *et al*. (2024b). (B) Simplified model of the Yang cycle and ethylene biosynthesis pathway, updated from Monthony et al. (2024) to reflect gene presence in *C. sativa*. (C) Updated model of the canonical ethylene signaling pathway, including ER-localized receptors, CTR1-mediated EIN2 phosphorylation, and EIN2-C–dependent regulation of ethylene responses via EBF1/2, EIN3/EILs, ERF1, and FYFs. Color indicates subcellular localization (ER-bound: blue, cytosolic: orange, nuclear: light orange); arrowheads denote activation (pointed) or inhibition (blunt), in the presence (green) or absence (red) of ethylene. Dashed arrows indicating indirect or abbreviated steps. Single, grey dashed arrow indicates movement. A detailed review of ethylene biosynthesis and signaling pathways can be found in Binder (2020).

The newly available fully phased XY genome allowed us, for the first time, to report putative ERGs present in both the X and Y chromosomes, including those localized in the pseudoautosomal region (PAR) and the non-recombining sex-determination region (SDR; Figure 2A). Ortholog analysis identified six X-specific ERGs, all within the putative SDR region of the X chromosome. Additionally, four ERGs were detected in the X chromosome PAR, with homologs in the Y chromosome PAR. Notably, no Y-specific ERGs were identified. A complete list of genes present in the X and Y haplotypes of the Otto II genome can be found in Supplementary Table 11. Compared to our earlier analysis using the older cs10 genome assembly, which identified a single *MTN* gene, the improved ortholog resolution enabled by the Pink Pepper and Otto II genomes revealed two MTN-family genes. These were provisionally named *CsMTN* and *CsMTN-LIKE1*, based on sequence similarity to *Arabidopsis MTN1* and *MTN2*. *CsMTN* matched both *MTN1* and *MTN2*, while *CsMTN-LIKE1* showed weaker similarity to *MTN2* and had been annotated as a nucleosidase-like gene in the *C. sativa* reference genome (NCBI 2023; Supplementary Table 11). Taken together, these new ERGs were integrated into an updated model for the Yang cycle, ethylene biosynthesis and signaling (Figure 2 B &C).

### Low nucleotide diversity suggests ERGs are highly conserved in *C. sativa*

To explore genetic variation within key CsERGs, we characterized single nucleotide polymorphisms (SNPs), structural variants (SVs), and nucleotide diversity (*θ_π_*) across the whole genome and 47 CsERGs. Variant calling on the genomes of XX and XY Deadly Kernel, La Rosca, and Panama Pupil V4 (n = 6 genomes) identified approximately 6.7 million polymorphic SNPs and ∼78 thousand SVs relative to the *C. sativa* ‘Pink Pepper’ reference genome. Within ±5 kb regions of the 47 CsERGs, 8,670 SNPs and 98 SVs were detected (Supplemental Figure 3A). Of these, 3,611 SNPs and 29 SVs were located within CsERGs gene sequences and were further examined for potential functional impact. The majority of variants were found in intronic regions or the 3’ and 5’ UTRs, and were classified as having modifier impact (Supplemental Figure 3B&C). Among coding region variants, most SNPs were synonymous, although 114 caused missense mutations, and 13 high-impact variants (7 SNPs and 6 SVs) were identified (Supplemental Figure 3 D&E). Notably, LOC115719259 (*CsMAT2*), LOC115716986 (*CsFYF3*), and LOC115717309 (*CsFYF4*) exhibited a high density of SNPs, primarily within introns, consistent with their predominantly (>90%) intronic gene structure. Additionally, LOC115706236 (*CsCTR1-LIKE*) and LOC115702329 (*CsETR1*) also showed elevated SNP counts (>100 SNPs). Across these five highly polymorphic genes, the average nucleotide diversity was 7.21 × 10⁻³, notably higher than the genome-wide average of 3.60 × 10⁻³ (Supplementary Table 12).

Across all CsERGs, the average nucleotide diversity was 2.85 × 10⁻³, indicating lower diversity than the genome-wide estimate. This signal was even more pronounced when excluding the five highly polymorphic genes, with the remaining 42 CsERGs exhibiting an average nucleotide diversity of 2.33 × 10⁻³. Interestingly, CsERGs located in sex chromosome subregions also displayed reduced variation: X-linked CsERGs within the SDR had an average nucleotide diversity of 1.77 × 10⁻³, while those located in the pseudoautosomal region (PAR) showed the lowest diversity overall at 1.46 × 10⁻³. The elevated SNP density observed in the five high-diversity genes may reflect underlying structural variants miscategorized during alignment.

### ACC Accumulation Differs by Sex and Stage

Endogenous ACC levels were quantified in *C. sativa* leaves at vegetative (day 0) and flowering (day 1) stages across experimental groups (Supplemental Figure 4A and Supplementary Table 13). Estimated marginal means from the linear mixed-effects model revealed that ACC concentrations were significantly higher in XY Day 1 - IFF compared to XX Day 0 - FF and XX Day 1 - FF (Tukey’s HSD, α = 0.05). No other groups displayed significant differences in ACC levels. No significant change in ACC concentration was observed between pre- and post-flowering XY plants, regardless of treatment. Additionally, IMF exhibited intermediate ACC levels between male (XY) and female (XX) floral phenotypes. SAM quantification was below the level of quantification and is presented as presence/absence in Supplementary Table 13.

Among all ERGs analyzed, *CsCTR1-LIKE* expression exhibited a strong and significant negative correlation with ACC concentration in XX Day 0 - FF (ρ = −0.8, p ≤ 0.001). However, no significant correlations were observed in other groups (Supplemental Figure 4B). Notably, all XY treatments exhibited weak positive correlations with *CsCTR1-LIKE* expression, suggesting a difference between *CsCTR1-LIKE* expression in XX and XY plants. Following STS treatment, IMF correlation remained strongly negative, while the control XX female flower exhibited only a weak correlation (Supplemental Figure 4B).

### Transcriptomics and Network analysis

#### Time and sex-driven transcriptomic patterns in XX and XY plasticity

To understand the effect of sexual plasticity treatments on ERG expression over time and tissue-type, transcriptomes were obtained from leaves (day 0 and day 1) and immature flowers (day 14) across treated, and untreated XX and XY plants of the DK, LR and PP genotypes. This yielded 139 transcriptomes, comprising ∼8 billion mapped reads with 24x coverage and high mapping quality (score = 103, insert size = 302 bp, Supplementary Table 8). PCA analysis found that the proportion of variance explained by the first two principal components increases over time (Figure 3A-C). At Day 0, PC1 and PC2 together account for 46.25% of the total variance (PC1: 29.07%, PC2: 17.18%). By Day 1, the combined variance explained increased to 61.94% (PC1: 41.66%, PC2: 20.28%), and at Day 14, it reached 74.05% (PC1: 49.41%, PC2: 24.64%). At day 0, PCA of ERG expression in untreated leaves showed a clear separation between XX and XY samples along PC1 (Figure 3A). The corresponding heatmap (Figure 3D) suggests that this clustering is driven by a subset of highly expressed ERGs in XX plants, including *CsCTR1/CTR1-LIKE*, *CsMTK1*, *CsFYF4/AGL42* and *CsACO3*.

**Figure 3.**
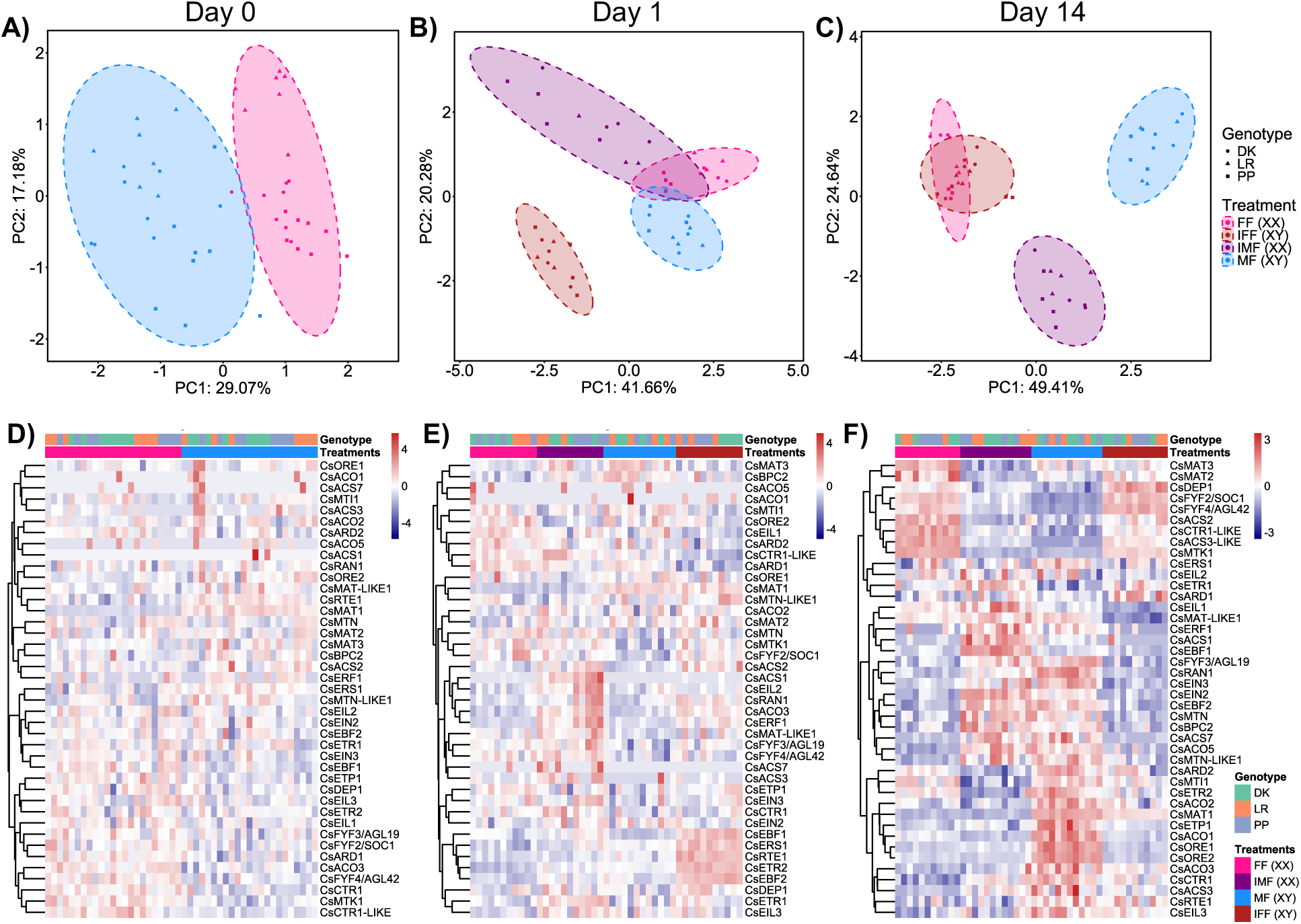
Principal Component Analysis (PCA) and heatmaps of 47 ERG expression across experimental time points and treatments. PCA was performed on batch-corrected variance-stabilized transformed (VST) counts of 47 ERGs. (A) Day 0 (pre-treatment, leaves); (B) Day 1 (post-treatment, leaves); (C) Day 14 (post-treatment, immature flowers). Each point represents an individual sample, and ellipses indicate the 95% confidence intervals for each group. Colors correspond to treatment groups, while symbols indicate cultivars (DK: Deadly Kernel, LR: La Rosca; PP: Panama Pupil V4). D-F) Heatmaps displaying expression patterns of 47 ERGs for each time point. Heatmap colors represent z-scores of gene expression, with rows corresponding to genes and columns to samples. The hierarchical clustering of genes (y-axis) was performed using Euclidean distance and complete linkage.

Following photoperiod change and application of the sex-altering treatments, changes in ERG expression patterns were observed at day 1. An increased variance is observed among treated groups, notably ethephon-treated XY plants (IFF) which showed a sharp divergence from other treatments along both principal components (Figure 3B). The heatmap for day 1 (Figure 3E) highlights elevated expression of ethylene signaling genes in IFF plants, including *CsEBF1*, *CsERS1*, *CsRTE1*, *CsETR2* and *CsEBF2*.

By day 14, PCA of ERG expression in immature flowers reveals a partial overlap between IFFs and the FF controls tissues (Figure 3C). The heatmap (Figure 3F) shows two distinct blocks of floral organ concordant (FOC) gene expression –shared gene expression among flowers of the same phenotypic sex. A female-biased, FOC block of ERGs including genes from the Yang Cycle (*CsMAT2/3*, *CsMTK1* and *CsDEP1*), ethylene biosynthesis (*CsACS2* and *CsACS3-LIKE*) as well as ethylene signaling and signaling-adjacent genes (*CsFYF2, CsFYF4* and *CsCTR1-LIKE*) are clustered in the top rows (Figure 3F).

In contrast, male expression patterns were more complex. PCA indicated that MFs and IMFs remained distinct, with separation along both principal components (Figure 3C). The heatmap (Figure 3F) revealed both FOC gene expression and the presence of uniquely expressed ERGs (uERGs) in MFs. Male-biased ERGs included *CsMTN/MTN-LIKE1* (Yang Cycle), *CsACS1*, *CsACS7*, *CsACO5* (biosynthesis), and multiple ethylene signaling-related genes (*CsEBF1, CsEBF2, CsERF1, CsFYF3, CsRAN1, CsEIN3, CsEIN2, CsBPC2*). Additionally, a set of uERGs was highly expressed only in MFs, including *CsARD2, CsMTI1, CsMAT1* (Yang Cycle), *CsACO2, CsACO1* (biosynthesis), and *CsETP1, CsETR2, CsORE1, CsORE2* (signaling).

#### Network analyses reveal XX and XY-unique ERGs involved in sexual plasticity

The PCA suggesting distinct XX and XY ERG expression patterns in response to sexual plasticity treatments led us to construct two separate sex-specific gene co-expression networks. This allowed for the identification of modules specific to each genotypic sex without cross-sex effects influencing module detection. WGCNA network construction for XX and XY plants resulted in networks with 12 and 16 module Eigengenes (ME), respectively. Hierarchical clustering dendrograms (Supplemental Figure 5A & B) display the gene co-expression relationships within these sex-specific networks. Distinct clustering patterns and variations in module compositions, size and number were observed between the XX and XY networks.

Module-trait correlation analyses were performed independently for the XX and XY networks to identify modules significantly associated with treatment and specific time points (Supplemental Figure 5C & D). Priority was given to modules strongly correlated with treatment to identify gene clusters most responsive to the experimental conditions. In the XX network (Supplemental Figure 5C), five modules displayed significant correlations with treatment: MEbrown (r = −0.89), MEblack (r = −0.52), MEmagenta (r = 0.54), MEblue (r = 0.66) and MEred (r = 0.71). For the XY network (Supplemental Figure 5D), three modules surpassed the threshold for significant correlation with treatment: MEgrey (r = 0.53), MEpink (r = 0.65), and MEgreenyellow (r = 0.5).

From these treatment-associated modules, ERGs were extracted for further investigation, identifying 19 ERGs in the XX network, and 6 ERGs in the XY network, resulting in a total of 24 unique ERGs (Supplementary Table 13 and Supplementary Table 14). To evaluate their functional significance within each module, intramodular connectivity (kME) was calculated. The most highly connected gene in the XX network was *CsACO5* (kME = 0.96), while in the XY network, *CsERF1* exhibited the highest connectivity (kME = 0.93). Interestingly, *CsERF1* was the only gene present in both networks, with a kME of 0.95 in the XX network. All 24 ERGs were retained for subsequent differential gene expression analysis.

#### Differential expression analysis reveals 14 candidate sexual plasticity ERGs

To refine the candidate ERGs involved in sexual plasticity, we conducted differential gene expression (DGE) analysis on the 24 ERGs identified from treatment correlated XX and XY WGCNA modules. Contrasts were set between treated and untreated samples of the same genotypic sex at time points following sexual plasticity treatment (Day 1 and Day 14), and genes were considered significantly differentially expressed if |log₂(Fold Change)| ≥ 1 and adjusted p-value ≤ 0.05 across all three genotypes. Of the initial 24 ERGs, 14 met these filtering criteria (grey highlighted genes in Supplemental Figure 2). The resulting refinements yielded a candidate ERG list including those most likely to play a role in ethylene-mediated sexual plasticity in *C. sativa* (Table 2).

**Table 2.**
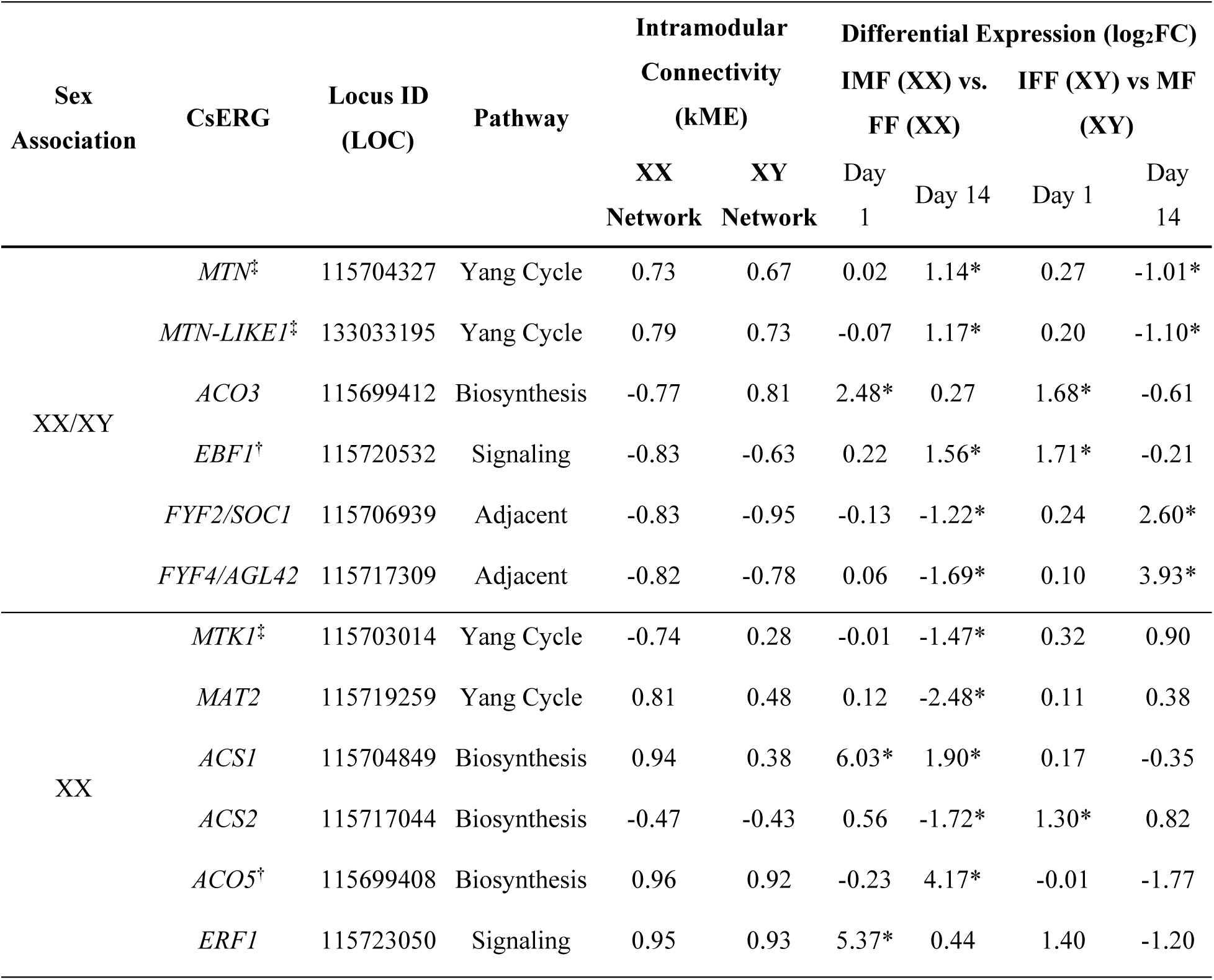
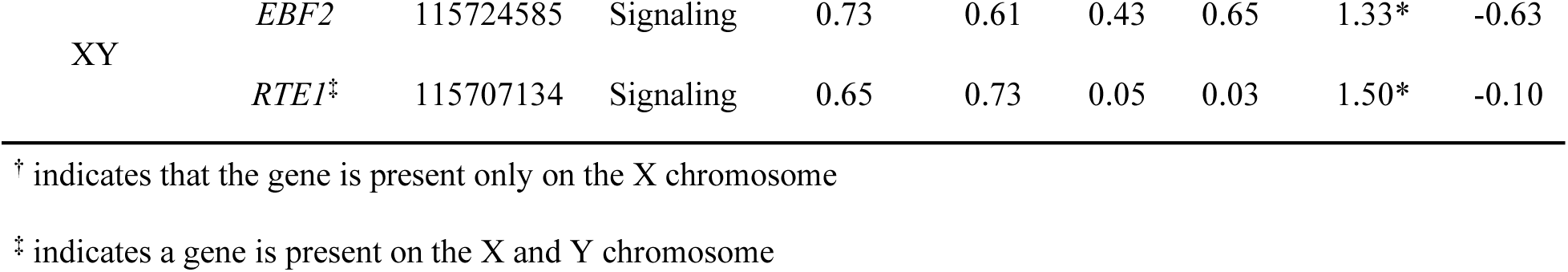
Candidate sexual plasticity ERGs in XX and XY *C. sativa*. Candidate ERG filtration included those identified with a significant WGCNA correlation between module and treatment (r ≥ 0.5; p-adjust ≤ 0.05) followed by a filtration using differential gene expression (DGE). DGE was conducted intra-sex on day 1 or day 14 retaining only those genes which were significantly (p-adjust ≤ 0.05) differentially expressed across all 3 tested genotypes. Log_2_FC values denoted by a * meeting filtration criteria for significant differential expression (p-adjust ≤ 0.05; |log_2_FC ≥ 1|; in 3 genotypes).

From the 14 candidate ERGs, six met the criteria for association with sexual plasticity in both XX and XY sex changed plants, a further six were associated with XX plants only and two with XY plants (Table 2). Within the XX and XX/XY-associated ERGs, there was representation from all pathways including: Yang cycle (e.g., *MTN*, *MAT2*), ethylene biosynthesis (e.g., *ACS1*, *ACO5*), signaling (e.g., *EBF1*, *ERF1*), and signaling adjacent regulatory pathways (e.g., *FYF4/AGL42*). In contrast, the XY-associated ERGs were primarily limited to signaling components (*EBF2*, and *RTE1*), perhaps reflecting the signaling response from the ethephon-induced sex change.

To identify temporal patterns underlying ethylene-induced sexual plasticity, we examined the differential expression of ethylene-related genes (ERGs) at two stages: in mature leaves immediately following treatment and photoperiod change (day 1) and in immature inflorescences during active sexual phenotype transition (day 14). Genes strongly associated with day 1 by WGCNA and/or differential expression analysis were classified as plasticity initiators, while those associated with day 14 were classified as plasticity stabilizers (Figure 4).

**Figure 4.**
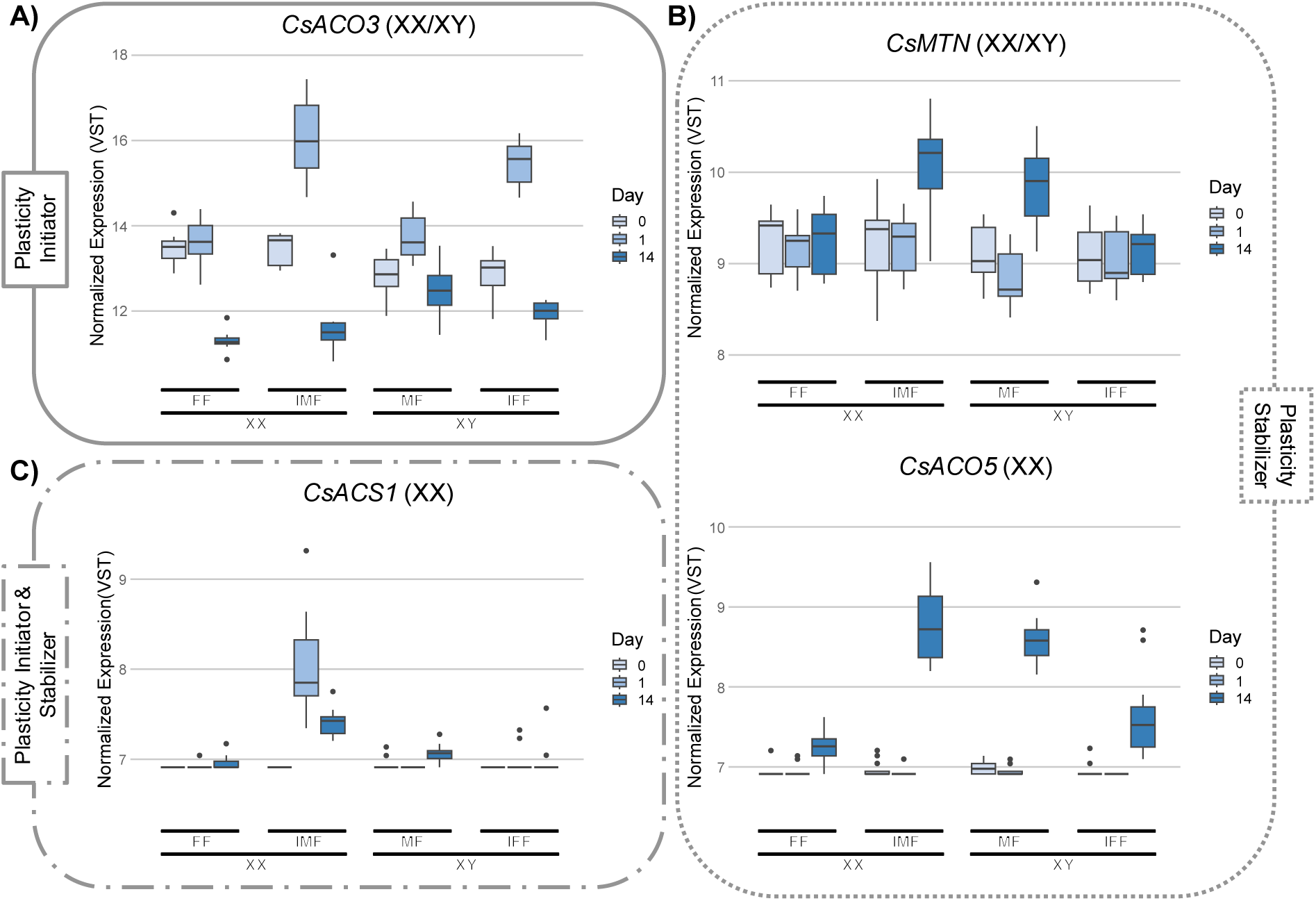
Representative VST normalized ERG RNA expression profiles for sexual plasticity initiators (A & C) and plasticity stabilizer genes (B), as identified using WGCNA and DGE filtration. Brackets next to names indicate chromosomal sex of plants where a significant effect of the gene on plasticity was identified.

In XX plants, three differentially expressed ERGs were significantly associated with plasticity initiation and ten associated plasticity stabilization (Table 2). Initiators included *ERF1* (log₂FC = 5.37) and *ACO3* (log₂FC = 2.48; Figure 4A), which were significantly upregulated in IMFs compared to FFs at day 1, suggesting early activation of ethylene biosynthesis and signaling pathways. By day 14, several ERGs displayed substantial differential expression, including strong upregulation of *ACO5* (log₂FC = 4.17; Figure 4B) and *EBF1* (log₂FC = 1.56), as well as moderate upregulation of the Yang cycle-associated genes *MTN* (log₂FC = 1.14; Figure 4B) and *MTN-LIKE1* (log₂FC = 1.17). In contrast, *MAT2* exhibited marked downregulation (log₂FC = –2.48). *ACS1* was the only ERG significantly differentially expressed at both timepoints in XX plants, with strong upregulation at day 1 (log₂FC = 6.03) and sustained, though reduced, expression at day 14 (log₂FC = 1.90), suggesting a role in both the initiation and stabilization of sexual plasticity (Figure 4C).

In XY plants, five ERGs were classified as initiators and four as stabilizers (Table 2). Two initiators, *EBF2* (log₂FC = 1.33) and *RTE1* (log₂FC = 1.50), were differentially expressed only in masculinized XY plants and did not show corresponding changes in XX plants. Among stabilizers, *FYF2* and *FYF4* were strongly upregulated during the phenotypic transition observed at day 14 (log₂FC = 2.60 and 3.93, respectively). *MTN* and *MTN-LIKE1*, which were upregulated in XX plants, were downregulated in XY plants (log₂FC = –1.01 and –1.10, respectively; Figure 4B).

The candidate ERGs were used to create a working model for the temporal control of ethylene-regulated sexual plasticity (Figure 5). This model incorporates plasticity initiators and plasticity stabilizers, highlighting their dynamic changes in sex- and treatment-specific expression profiles over time. Importantly, candidate ERGs critical to sexual plasticity regulation appear consistently at pathway bottlenecks, where alternate signaling routes are absent or limited. Key examples include *MTN* and *MTK1* at the initiation of the Yang cycle, *ACS1/2* and *ACO3/5* in the biosynthesis pathway (Figure 2B & Figure 5), and *EBF1/2* and *ERF1* in downstream signaling (Figure 2C & Figure 5).

**Figure 5.**
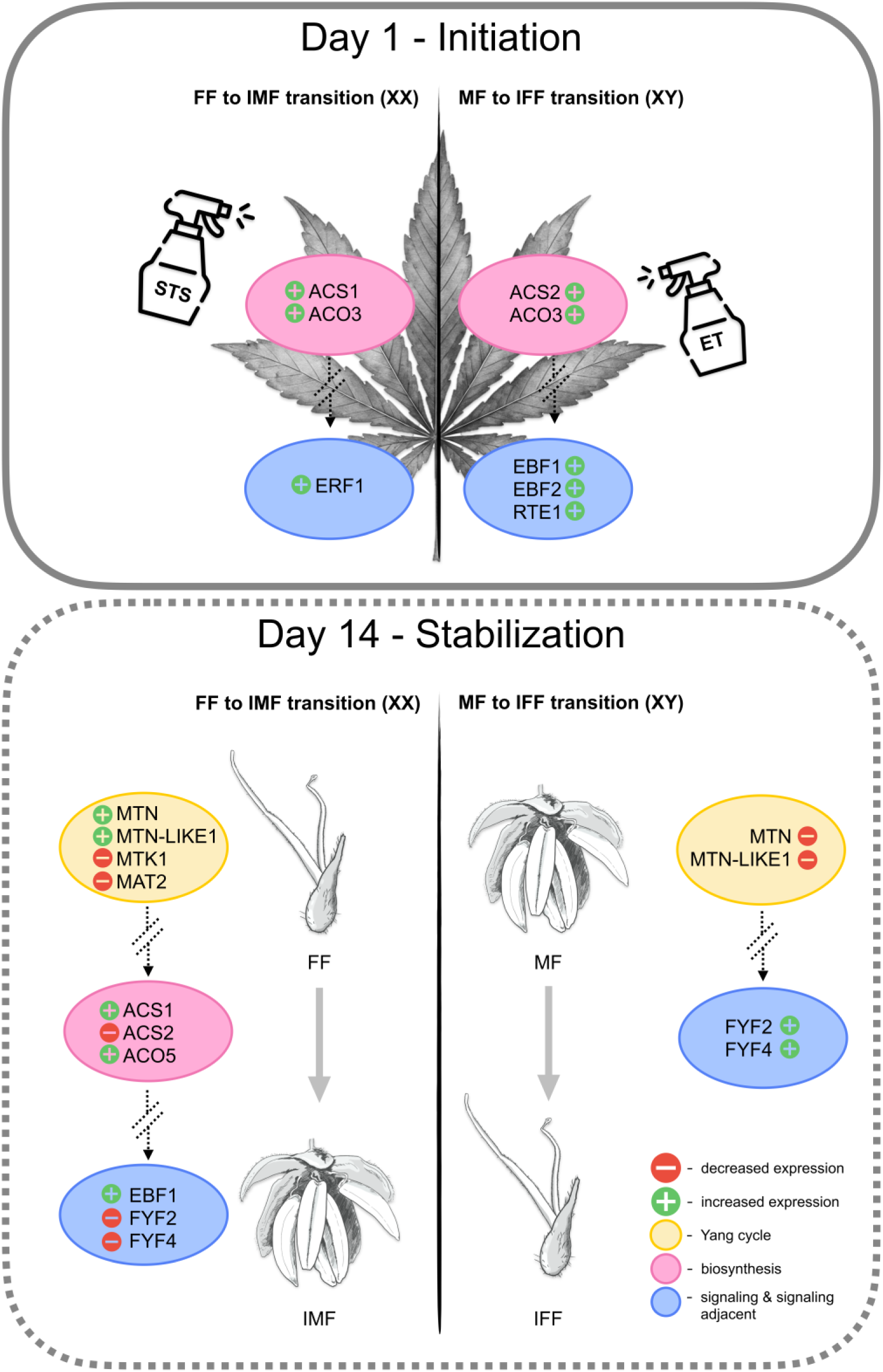
Working model of ethylene-induced sexual plasticity in *C. sativa*. The model outlines two phases: initiation, marked by broad upregulation of ethylene biosynthesis and signaling genes in both IMFs (XX) and IFFs (XY); and stabilization, when ERG expression diverges between IMF and IFF plants. Key expression changes span the Yang cycle, biosynthesis, signaling, and adjacent pathways, and are shown as directional symbols (green “+” for upregulation, red “–” for downregulation) based on log₂FC relative to untreated controls. FF→IMF and MF→IFF transitions reflect contrasts with untreated FFs and MFs, respectively. Flower illustrations by Alexandra K. McGregor.

## Discussion

### Ethylene-induced sexual plasticity is a dynamic, sex-specific process

Sexual plasticity, the capacity of a plant to shift its phenotypic sex from its baseline genetic sex in response to internal or external cues, is a phenomenon observed in many monoecious and dioecious species (Lin et al. 2016; Cossard and Pannell 2021; Käfer et al. 2022). In *Cannabis sativa*, there is long-standing experimental and anecdotal evidence supporting ethylene as a central regulator of this process (Galoch 1978; Moon et al. 2020; Adal et al. 2021). In this study, application of sex-altering treatments (silver thiosulfate for XX plants and ethephon for XY plants) reliably triggered phenotypic sex change in >80% of treated plants, confirming a conserved, inducible plasticity response (Table 1), achieving inflorescence sex change rates comparable to previous reports (Lubell & Brand, 2018; Moon et al., 2020).

Despite these shared phenotypic outcomes, our results reveal that XX and XY plants respond to ethylene modulation through distinct transcriptional programs. Analysis of 47 ethylene-related genes (ERGs) across time and tissue type uncovered a temporally dynamic, sex-specific response pattern, supporting a two-phase model of plasticity: (1) an early initiation phase, where shared upregulation of biosynthesis genes occurs shortly after treatment alongside divergent patterns of expression of ethylene signaling genes, and (2) a later stabilization phase, in which ERG expression diverges and consolidates new phenotypic sex identities (Figure 3; Figure 5). In the following sections, we explore how sex chromosome structure and gene location shape these responses, and how specific ERGs act as candidate regulators of each plasticity phase.

### Plasticity initiation: ERG responses shortly after sex-change induction

Ethylene biosynthesis and signaling are the two gene families that make up the early responsive ERGs. Of particular interest are the biosynthesis genes *CsACS1* and *CsACS2*, which encode ACC synthase, and *CsACO3*, which encodes ACC oxidase—together catalyzing the two-step conversion of SAM to ethylene (Binder 2020). *CsACS1* was strongly upregulated in STS-treated XX plants (IMFs) at Day 1 (log₂FC = 6.03) and remained elevated at Day 14 (log₂FC = 1.90), indicating a potential dual role in initiating and stabilizing masculinization (Figure 5 & Table 2). Normalized expression data showed a sharp spike in *CsACS1* expression at Day 1 in IMFs (Figure 4C), accompanied by a modest increase in ACC concentration (Supplemental Figure 4), supporting its role as a plasticity-associated ethylene biosynthesis gene. Across all other tissues and treatments, including MFs and IFFs, *CsACS1* remained minimally expressed. In contrast, *CsACS2* displayed a broader and less clearly sex- or treatment-specific pattern. In XY plants, *CsACS2* was upregulated following ethephon treatment at Day 1 (log₂FC = 1.30) and downregulated in IMFs by Day 14 (log₂FC = –1.72), reaching levels similar to MFs. However, the magnitude of these changes was comparatively small. Normalized expression showed that *CsACS2* expression was distributed across multiple tissue types and timepoints, including elevated expression in IFFs and Day 1 MFs—suggesting a more constitutive role in ethylene production. Previous studies have reported similarly variable *CsACS2* expression. Our previous study of ERGs found higher *CsACS2* expression in FFs compared with IMFs and MFs (Monthony et al. 2024), while Borin *et al*. (2024) found no significant expression differences across FF, MF, and IMF flower tissues at Day 14 using qRT-PCR. These conflicting results may reflect tissue-type, developmental stage, or genotype-specific variation rather than a direct role in sexual plasticity. In other plant species, such as cucurbits and tomato, members of the ACS gene family show tissue- and stage-specific expression, with functionally distinct roles despite similar enzymatic activity (Van de Poel et al. 2014; Boualem et al. 2015). *CsACS2* may follow a similar pattern in *C. sativa*, highlighting the need for finer-resolution temporal and tissue-specific analyses to clarify its function.

*CsACO3* was also differentially expressed in leaf tissues one day after sex-change induction, showing significant upregulation in both IMFs and IFFs (log₂FC = 2.48 and 1.68, respectively; Table 2). Although the magnitude of change was greater in XX plants, the consistent direction across both sexes suggests *CsACO3* may serve as a shared ethylene biosynthesis initiator during early reprogramming. This pattern aligns with its expected enzymatic function and may reflect a transient upregulation to support immediate ethylene flux following hormonal induction. Further exploration of its expression in floral tissues and later timepoints would help determine whether *CsACO3* has a continued or transient role in sexual plasticity. Interestingly, our previous study of ERGs in *C. sativa* sexual plasticity using flower tissues from MF, FF, and IMF plants (Monthony et al. 2024) found that *CsACO3* expression was highest in MFs and lowest in FFs and IMFs, suggesting a karyotype-concordant pattern of expression. In the current dataset, we again observe highest expression in MFs at Day 14, with IFFs showing comparable levels—further validating this karyotype-linked trend (Figure 4A). This study newly implicates *CsACO3* in early reprogramming, underscoring its dynamic, stage-specific expression.

Despite the sharp transcriptomic induction of *CsACS1* and *CsACS2* following sex-change treatments, this response was not fully mirrored in the measured ACC concentrations. For example, despite substantial upregulation of *CsACS1* (log₂FC = 6.03) in leaves of the IMF treatment relative to FFs, the ACC levels remained similar between these groups (Supplemental Figure 4). In contrast, IFFs exhibited the highest ACC concentration of all groups, despite a more modest *CsACS2* induction (log₂FC = 1.30), which more closely aligns with the anticipated relationship between gene induction and metabolite output. This apparent disconnect between *ACS* expression and *ACC* accumulation is consistent with previous findings in tomato, where tissues producing the most ethylene—such as the pericarp—had relatively low ACC content and ACS activity, while abundant *ACC* transcript levels were observed in tissues with limited ethylene production (Van de Poel et al. 2014). These discrepancies have been interpreted as evidence for tight spatial and temporal regulation of ACC metabolism and transport, as well as the rapid conversion of ACC to ethylene in actively ripening or differentiating tissues. Possible explanations for the elevated ACC levels observed in IFFs may reflect a lag between biosynthetic induction and ACC turnover, a localized ethylene amplification loop triggered by exogenous ethephon, or even the accumulation of ACC as a metabolic reserve—potentially stockpiled for future ethylene biosynthesis depending on tissue-specific developmental cues. Together, these findings highlight that ACC concentrations alone are insufficient indicators of ethylene biosynthetic activity, particularly during rapid transcriptional reprogramming. This underscores the complex interplay of transcriptional regulation, enzyme kinetics, and tissue-specific flux. Clarifying the roles of *CsACS1* and *CsACS2* in ethylene production will likely require future studies incorporating spatial transcriptomics and enzyme activity assays.

*ERF1*, a central transcriptional activator in the ethylene signaling pathway, acts downstream of *EIN3* and *EIL1* to mediate broad ethylene responses through direct activation of GCC-box-containing target genes (Solano et al. 1998; Binder 2020). Given the well documented role of ethylene response factors (ERFs) in ethylene-mediated developmental programming, and the recent association of *CsERF106* with sexual plasticity (Orozco et al. 2025), *CsERF1* is a plausible candidate for ethylene mediated sexual plasticity. In our dataset, *CsERF1* was strongly upregulated in STS-treated XX plants at Day 1 (IMFs vs FFs; log₂FC = 5.37) and exhibited the highest kME values of all candidate genes in both the XX (0.95) and XY (0.93) WGCNA networks, indicating tight integration with other genes responsive to sex change treatments. Although the IFF vs MF contrast in XY plants showed a log₂FC of 1.40, this effect did not meet our criteria for statistical significance due to genotype-level variability (Supplemental Figure 2). Nevertheless, this trend suggests that *CsERF1* may be responsive to ethylene exposure across both sexes, with a potentially conserved role in reprogramming floral identity. Taken together, these findings support a role for *CsERF1* as an early transcriptional integrator in ethylene-mediated sexual plasticity, particularly in XX plants, with evidence for broader involvement that warrants further investigation.

### Plasticity stabilization: Genes consolidating new sexual identity

*CsACO5*, an X-linked gene with no Y copy, emerged as a compelling candidate for the stabilization of male floral identity during sexual plasticity, based on its strong and consistent upregulation in IMFs. Despite being X-linked, *CsACO5* levels are nearly identical in XY MFs and XX IMFs by Day 14, implying regulation by the male-flower program, not chromosome count (Figure 4B). Similarly, *CsACO5* expression in IFFs resembled that of FFs, with both groups exhibiting lower transcript levels. This floral organ concordant (FOC) expression pattern—where gene expression reflects phenotypic sex rather than chromosomal sex—is consistent with Borin *et al*. (2024) and Monthony *et al*. (2024), both of whom reported elevated *CsACO5* expression in IMFs compared to FFs. At Day 14, *CsACO5* expression was markedly upregulated in IMFs relative to untreated FFs (log₂FC = 4.17), while normalized expression in MFs was comparable to IMFs, and IFFs exhibited reduced expression levels similar to FFs, though with genotype-dependent variability (Supplemental Figure 2E). *CsACO5* remains low at Day 0 and Day 1 in all groups (Figure 4B).

Complementing this late-stage induction, another gene that may contribute to the stabilization of male floral identity is *CsACS1*, which showed strong early induction in STS-treated XX plants (IMFs; log₂FC = 6.03 at Day 1) and remained significantly upregulated at Day 14 (log₂FC = 1.90), suggesting a role in both initiating and stabilizing masculinization (Table 2 & Figure 5). Normalized data showed a sharp spike in *CsACS1* transcript abundance at Day 1 in IMFs (Figure 4C), accompanied by a modest increase in ACC concentration (Supplemental Figure 4), supporting its role as a plasticity-associated ethylene biosynthesis gene. *CsACS1* remained minimally expressed across all other tissues and treatments, including MFs and IFFs.

Comparative studies in other ethylene-sensitive species provide context for these dynamics. Studies in cucumber and melon have shown that disruption of specific *ACS* and *ACO* genes can alter floral sex ratios and reduce ethylene production, highlighting their context-dependent roles in sex expression. In cannabis, *CsACS1* and *CsACO5* may act as stabilizers, with *CsACS1* showing early and persistent upregulation, and *CsACO5* induced specifically at Day 14, together promoting male floral development or suppressing female identity. Floral ethylene dynamics are also linked to developmental timing, with genes delaying senescence indirectly modulating ethylene output. For example, predicted orthologs of Arabidopsis *FOREVER YOUNG FLOWER* (*FYF*), *CsFYF2* (LOC115706939) and *CsFYF4* (LOC115717309), were consistently upregulated in phenotypically female flowers (FFs and IFFs). FYF transcription factors are known to repress ethylene-mediated senescence and abscission pathways, extending floral longevity in early developmental stages (Chen et al. 2021). Similarly, Shi *et al*. (2024a) reported elevated *CsFYF2*—LOC115706939, annotated as *CsSOC1*—in female meristems and flowers, complementing our observation of significantly lower *CsFYF2/4* expression in MFs and IMFs (Table 2). These expression trends align with the longevity role of *FYF* proposed in other species, and the shorter developmental lifespan of male cannabis inflorescences compared to female ones (Shi et al., 2024a). These findings support a model in which *CsACO5* and *CsACS1* stabilize male floral identity during inflorescence development, while *CsFYF2* and *CsFYF4* modulate sex-specific senescence trajectories in a phenotype-dependent manner.

Upstream of both *ACS* and *ACO* activity, the Yang cycle contributes essential precursors for ethylene biosynthesis. One gene in this pathway, *CsMAT2*, an autosomal Yang cycle gene located on chromosome 2, encodes methionine adenosyltransferase (MAT or SAMS), which synthesizes S-adenosylmethionine (SAM)—the first substrate used by ACS in ethylene biosynthesis (Figure 2A). *CsMAT2* was significantly downregulated in IMFs compared to FFs at Day 14 (log₂FC = – 2.48), a developmental window coinciding with early pollen maturation (DiMatteo et al. 2020 & Figure 1), suggesting a potential role in male gametophyte development. This is supported by findings in Arabidopsis and rice, where *MAT* gene disruption impairs pollen tube growth, male fertility, and seed set (Chen et al. 2013, 2016b). This is especially relevant in *C. sativa*, where STS-induced male flowers often exhibit poor pollen quality (Lubell and Brand 2018; DiMatteo et al. 2020). Notably, in pumpkin, a long non-coding RNA has been shown to stabilize SAMS protein and promote fruit development via ethylene biosynthesis (Tian et al. 2024), illustrating how *MAT* regulation can directly modulate ethylene-dependent outcomes.

Beyond its potential functional relevance, *CsMAT2* also shows interesting network and evolutionary characteristics. *CsMAT2* had strong connectivity in the XX network (kME = 0.81) but weaker integration in XY plants (kME = 0.48). It also had the highest nucleotide diversity among ERGs (π = 5.99 × 10⁻³), exceeding the genome-wide average (Supplementary Table 12). This may reflect relaxed selection on *CsMAT2* due to redundancy in the *MAT* gene family as four *MAT/SAMS* genes are present in the *C. sativa* genome (Figure 2B). A similar pattern occurs in tomato, where distinct *SlMAT* paralogs exhibit tissue- and stage-specific expression during ethylene-mediated fruit ripening (van de Poel et al. 2012). These findings suggest that *CsMAT2* may represent a functionally specialized member of the *MAT* family in cannabis, with a potential role in early floral development that merits further investigation.

### Genomic context of plasticity: ERG distribution and chromosomal constraints

The spatial distribution of ERGs across the genome provides insights into the genetic architecture underlying sexual plasticity. Of the 47 ERGs identified, 12 were sex-linked: six in the pseudoautosomal region (PAR) and six within the non-recombining SDR of the X chromosome (Figure 2A), with none in the Y-linked SDR. Although suppressed recombination typically reduces nucleotide diversity in SDR regions relative to the PAR, we observed unexpectedly lower nucleotide diversity among PAR-located ERGs (θπ = 1.46 × 10⁻³) compared to SDR ERGs (θπ = 1.77 × 10⁻³). Both regions exhibited nucleotide diversity levels below the autosomal ERG average (θπ = 3.23 × 10⁻³), suggesting strong purifying selection on PAR-localized genes, particularly *CsMTK1*, *CsRTE1*, and *CsMTN*/*MTN*-*LIKE1*, which lack autosomal homologs in their respective enzymatic pathways. These genomic patterns highlight how chromosomal localization shape ERG function during sexual plasticity.

### ERGs in the sex-determining region (SDR)

All six X-specific ERGs within the SDR have homologs elsewhere in the genome (Figure 2A), functioning either in ethylene biosynthesis (*CsACS3-LIKE* and *CsACO5*) or signaling (*CsETR1*, *CsCTR1*, *CsEBF1*, and *CsEIL1*). *CsACO5*, previously discussed as a key stabilizer of male floral identity in XX plants, and *CsEBF1* are particularly noteworthy for their localization to the SDR. This genomic configuration is reminiscent of sex-determining regions in other dioecious species, such as kiwifruit (*Actinidia*), where lineage-specific duplication events gave rise to genes acquiring novel, sex-specific functions through regulatory divergence (Akagi et al. 2018). In cannabis, *CsACO5* and *CsEBF1*, emerged as candidate regulators of sexual plasticity (Table 2). *CsEBF1* and its autosomal counterpart, *CsEBF2*, encode F-box proteins that target the *EIN3/EIN3-LIKE* (*EIL1-3*) transcription factor family for degradation, acting as negative regulators of ethylene signaling in its absence (Binder 2020). Unlike the canonical Arabidopsis model where EBF mRNA is suppressed upon ethylene exposure (An et al. 2010), both cannabis homologs were unexpectedly upregulated in ethephon-treated IFFs at Day 1 (*EBF1* log₂FC = 1.71; *EBF2* log₂FC = 1.33). By Day 14, *CsEBF1* was also significantly upregulated in IMFs, a pattern not shared by *CsEBF2*. *CsEBF1* expression in sex changed flowers (day 14) suggests a potential role in stabilizing masculinization by suppressing ethylene signaling. As an SDR-linked gene, its regulation may be influenced by sex chromosome context, but further research is needed to determine whether this reflects sex-specific control or a more general role in floral development under STS treatment.

### ERGs in the pseudoautosomal region (PAR)

We identified six ERGs shared within the PAR of the X and Y chromosomes. Among them, *MTN,* which is essential for recycling methylthioadenosine (MTA), a byproduct of the first committed step in ethylene biosynthesis. *CsMTN* and *CsMTN-LIKE1* were located within a WGCNA module correlated with sexual plasticity treatment in XX plants (r = 0.66) and with high module membership (kME = 0.73 and 0.79, respectively), supporting their centrality in the transcriptional response to masculinization. Both genes were significantly upregulated in IMFs relative to untreated XX controls (log_2_FC = 1.14 and 1.17), while IFFs showed significant downregulation relative to XY controls (log_2_FC = −1.01 and −1.10; Table 2). These findings align with our previous study showing floral organ concordant expression of *MTN* (Monthony et al. 2024) and with Borin *et al*. (2024), who reported high *CsMTN* expression in MF and IMFs. In Arabidopsis, functional analyses have shown that *MTN* is required for male fertility: double mutants of *mtn1-1 mtn2-1* exhibit indehiscent anthers and malformed pollen grains (Waduwara-Jayabahu et al. 2012). In dioecious *Amborella trichopoda*, transcriptomic analyses of immature flowers revealed differential expression of *AmbMTN* by sex, with higher expression in female tissues (Carey et al. 2024a). Whether this response reflects direct ethylene regulation or secondary developmental reprogramming remains to be clarified, as Yang cycle gene responsiveness to ethylene varies across species (Bürstenbinder et al. 2007; van de Poel et al. 2012)

To further assess the role of methionine recycling during sexual plasticity, we next examined *CsMTK1*, a downstream Yang cycle enzyme also localized in the PAR (Figure 2A). *CsMTK1* phosphorylates 5-methylthioribose (MTR), a product of MTN activity, to form 5-methylthioribose-1-phosphate (Figure 2B). Only one *MTK* is present in *C. sativa*, consistent with *A. thaliana*, *Plantago major* L., and several other species (Pommerrenig et al. 2011; Chen et al. 2025a). Whereas *CsMTN* and *CsMTN-LIKE1* were strongly connected to plasticity modules in both XX and XY networks, *CsMTK1* expression was XX-specific (*kME* = –0.74) and weakly connected in XY plants (*kME* = 0.28), suggesting limited involvement in male-to-female transitions. *CsMTK1* was significantly downregulated in IMFs compared with FFs (log₂FC = – 1.47), with no significant change between IFFs and MFs. This Day 14-specific downregulation suggests *CsMTK1* is developmentally, rather than ethylene, regulated. This aligns with Arabidopsis and rice, where MTK is essential for SAM recycling under sulfur limitation but not induced by ethylene (Bürstenbinder et al. 2007; Rzewuski et al. 2007). By contrast, *MTK* expression is ethylene-responsive in climacteric species such as tomato and cucumber, pointing to a species-specific regulatory architecture (van de Poel et al. 2012; Chen et al. 2025a). In *Cannabis*, these patterns support a functional decoupling between *CsMTN* and *CsMTK1* during sexual plasticity, wherein MTA cleavage via MTN is prioritized while methionine recycling through MTK1 is developmentally downregulated. Supporting this, *CsMTK1* exhibited elevated nucleotide diversity (π = 4.58 × 10⁻³), higher than both the PAR ERG average and genome-wide mean (Supplementary Table 12), suggesting relaxed purifying selection relative to *CsMTN*. Together, the gene expression and diversity data suggest *CsMTK1* may be dispensable for ethylene-mediated sex change in *Cannabis*.

The final candidate ERG located in the PAR is *CsRTE1*, a known negative regulator of ethylene signaling. Unlike the Yang cycle genes discussed above, which may play a role in modulating the biosynthetic capacity of the pathway, *CsRTE1* acts in the ethylene signaling pathway, appearing as a responsive gene rather than a driver of sexual plasticity. In our study, *CsRTE1* was significantly upregulated in XY leaf tissue one day after ethephon treatment (Day 1 IFFs), a pattern not observed in treated XX plants. This response is consistent with previous findings in *Arabidopsis*, where *RTE1* is induced by ethylene and functions in a feedback loop to stabilize the *ETR1* receptor and attenuate downstream signaling (Resnick et al. 2006). The transient upregulation of *CsRTE1* in XY plants may reflect a mechanism for buffering early ethylene signaling—in this case in response to ethephon treatment—and preventing an overactivation of the autocatalytic ethylene biosynthesis loop. Supporting this, ACC concentrations in Day 1 IFFs were not significantly elevated compared to untreated XY plants (Supplemental Figure 4), suggesting that ethephon application does not trigger a runaway ethylene biosynthesis response within first 24 hours following ethephon treatment. Instead, *CsRTE1* may help temper responses to exogenous ethephon applications, maintaining signaling homeostasis as phenotypic feminization is initiated.

### Convergent phenotypes, divergent mechanisms: A phase-dependent model

The integration of temporal transcriptomic patterns, pathway-specific metabolite data (ACC), and ERG genomic localization underscores the modularity and adaptability of the ethylene response network in *Cannabis*. Despite a shared hormonal signaling pathway, XX and XY *Cannabis sativa* plants follow distinct molecular trajectories during sex change yet converge on similar phenotypic outcomes: the production of opposite-sex flowers. This apparent contradiction of convergent floral phenotypes arising from divergent gene expression programs, is a central finding of this study, and supports a refined, phase-dependent model of ethylene-regulated sexual plasticity in *C. sativa* (Figure 5).

In this model, plasticity begins prior to floral primordia emergence, shortly after STS or ethephon application and photoperiod shift. This initiation phase is marked by broad but sex-specific upregulation of ethylene biosynthesis and signaling genes. Shared early responders include *CsACO3* (upregulated in both XX and XY contexts), *CsACS1* and *CsERF1* in IMFs, and *CsACS2*, *CsRTE1*, and both *CsEBF1/2* in IFFs. These genes likely initiate developmental reprogramming by shifting the hormonal balance toward a new sex-specific pathway.

In the subsequent stabilization phase, distinct ERGs help maintain the new phenotypic identity. In masculinized XX plants, *CsACO5*, *CsMTN*, and *CsACS1* remain upregulated, supporting male floral development. In feminized XY plants, upregulation of *CsFYF2* and *CsFYF4* may suppress senescence and promote female organ longevity. These stabilization-phase genes often show sex-biased expression, with some (e.g., *CsACO5*, *CsEBF1*) located in the SDR and others (e.g., *CsMAT2*, *CsERF1*) on autosomes. These findings suggest that ethylene-mediated plasticity is not governed by a single sex-linked switch, but rather arises from distributed, tissue- and time-specific regulatory modules.

Our multi-omic analysis provides the first temporally resolved, genome-informed framework for ethylene-mediated sex change in *Cannabis sativa*. By capturing both conserved and sex-specific ERG responses across genotypes, we identify key plasticity initiators (e.g., *CsACO3*, *CsERF1*), stabilizers (e.g., *CsACO5*, *CsACS2, CsFYF2/4*), and dual-phase regulators (*CsACS1*), and propose a working model for understanding ERG function in sexual plasticity (Figure 5). The study also highlights candidate sex-specific regulators such as *CsMAT2*, *CsMTN*, and *CsMTK1* in XX contexts, and *CsRTE1* and *CsEBF2* in XY plants. Looking ahead, spatial transcriptomics and functional validation through gene silencing or editing will be critical to dissect causal roles within this regulatory network and advance the development of sex-stable genotypes for breeding and cultivation. As our understanding deepens, future models should also incorporate the likely crosstalk between ethylene and other plant growth regulators, such as gibberellins and cytokinins, which are known to influence sex expression in *Cannabis sativa*.

## Materials & Methods

### Plant selection and vegetative growing conditions

Vegetative mother and father plants from three commercially available drug-type genotypes—*La Rosca*, *Panama Pupil V4*, and *Deadly Kernel*—were greenhouse-grown, cloned, and rooted for two weeks under LED lighting (Phlizon, China) in moistened stone wool (Indoor Farmer, Canada) beneath vented plastic humidity domes. Rooted clones were then transferred to 10 cm square pots filled with sterilized Pro-Mix BX (Pro-Mix, Canada) and maintained under controlled environment conditions in growth chambers (Conviron, Canada) for 10 days to allow root establishment. During this period, plants were watered by hand as needed.

Subsequently, plants were transplanted into 4-litre round pots and grown for 12 days under a 12/12 day/night photoperiod with drip irrigation adjusted weekly. Thereafter, the photoperiod was shifted to 18/12 to induce flowering. Photosynthetic photon flux density (PPFD), as well as day and night humidity levels, were gradually decreased throughout the experiment, reaching a maximum of 750 μmol/m²/s at canopy height, as detailed in Supplementary Table 1. The pH of the growth substrate was monitored weekly and maintained between 5.5 and 6.0, adjusted as necessary using 1.5 g/L of elemental sulfur during watering. A complete description of growing conditions, including photoperiod, light intensity, and fertigation regimes, is provided in Supplementary Table 1.

### Plant sexing

Samples were taken from 6-month-old vegetative mother and father plants, in all three drug-type genotypes to confirm the chromosomal sex. Approximately 50 mg of young leaf tissue was harvested for DNA extraction and desiccated for four days using Drierite (Xenia, OH, USA). The dried tissue was ground using metallic beads in a RETSCH MM 400 mixer mill (Fisher Scientific, MA, USA) and DNA was extracted using a CTAB-chloroform protocol (Lapierre et al. 2023b). Briefly, powdered tissue was treated with CTAB buffer, followed by phenol-chloroform extraction. The resulting DNA pellet was washed with ethanol, resuspended in water, and quantified using a Qubit fluorometer with the dsDNA HS assay kit (Thermo Fisher Scientific, Waltham, MA, USA). DNA concentrations were adjusted to 10 ng/µL.

Sex determination was performed using PCR Allelic Competitive Extension (PACE, 3CR Bioscience Ltd) assay using CSP-2 primers (Supplementary Table 2) and methodology as described in Toth (2022), and Toth *et al*. (2020). PACE reactions were performed with PACE 2.0 Genotyping Master Mix (3CR Bioscience Ltd) and thermally cycled according to the manufacturer’s instructions, with five extra final cycles (i.e. a total of 35 cycles of denaturation, annealing and extension) on the Applied Biosystems Veriti Thermal Cycler. Post-PCR fluorescence signals were detected with the Applied Biosystems 7500 Real-Time PCR System and analyzed using Applied Biosystems 7500 Software v2.3.

### Plant flowering and sex plasticity induction

All plants were cultivated under LED lighting in controlled-environment growth chambers. To prevent pollination and accommodate space constraints, plants were grown in two separate batches under identical conditions (Supplementary Table 1). The first batch contained male flowering treatments: MF (XY) and IMF (XX), while the second batch contained female flowering treatments: FF (XX) and IFF (XY). Induction of sexual plasticity in XX plants was performed using STS, following the protocol of Jones & Monthony (2022). A 3 mM silver thiosulfate (STS) solution with 0.1% Tween 20 was prepared from sodium thiosulfate (Fisher Chemical, Fair Lawn, NJ, USA) and silver nitrate (Fisher Chemical, USA). The solution was applied as a foliar spray (∼50 mL per plant) once per week for three weeks, ensuring thorough leaf coverage. The first application coincided with the photoperiod shift, immediately after the first 12-hour night, and all subsequent treatments were administered immediately after growth chamber lights extinguished in order to prevent leaf burning.

Induction of sexual plasticity in XY plants was achieved using 500 mg/L ethephon, adapted from Moon et al. (2020). The treatment solution was prepared from a 40% aqueous solution of ethephon (2-chloroethylphosphonic acid; AK Scientific, Union City, CA, USA), by diluting it in distilled water to the final concentration (500 mg/L; 8.65 mM) with 0.1% Tween 20 as a surfactant. Plants were sprayed to saturation (∼50 mL per plant) in a single application, administered immediately after the photoperiod shift when growth chamber lights extinguished, to prevent leaf burning. In both batches, control plants (MF and FF) were sprayed to saturation with distilled water containing 0.1% Tween 20 to ensure comparable treatment conditions.

### Floral development and quantification of sexual plasticity

Floral development was tracked weekly for 28 days following floral induction. Representative floral clusters were selected from the top half of the plant for macroscopy. Plants were visualized using an Olympus SZX16 macroscope (Olympus Corp., Japan) and photographed with an Olympus DP21 camera. Sexual plasticity was quantified by determining the ratio of flowers on six randomly selected inflorescence segments measuring 3 cm whose phenotypic sex no longer matched the plants genotypic sex. This was done following 28 days of flowering. For control plants (MF and FF), which displayed no sexual plasticity, one randomly selected inflorescence segment was selected.

Floral sex ratios were calculated in R (v4.1.1). A two-way fixed effects analysis of variance (ANOVA) was conducted. Sex change (%) was the response variable, genotype and phenotypic floral sex were the independent variables. Prior to analysis of variance, the sex change data were transformed using an arcsine square root transformation (McDonald 2014), as the datasets for sex change (in percentiles) were highly clustered at 0 and 1, and the arcsine square root model shows less dramatic variance at the end of the distribution (Monthony et al. 2021a). Means comparisons was performed using a Tukey-Kramer post-hoc test (Kramer 1956) to account for multiple comparisons. Post-hoc means comparisons were visualized using the *multcompLetters4* from the multcompView package (Piepho 2004). Bar plots of the average sex change were generated with ggplot2 (v3.4.1; Wickham 2009) using the Paired colour palette from RColorBrewer (Brewer et al. 2002).

### Cannabinoid quantification

Cannabinoid quantification was performed using an established UPLC-PDA method, as previously described by Lapierre et al. (2023b), at the Metabolomics Platform, Institute of Nutrition and Functional Foods (INAF), Université Laval, Québec, QC, Canada. In brief, plants were grown under standard greenhouse conditions to maturity, as outlined in Lapierre et al. (2023b) and mature female (XX) flowers (∼5 g) were collected, trimmed and dried for analysis. A subsample (200 mg) was extracted in 25 mL of 80% methanol (Thermo Fisher Scientific, Optima™ HPLC ≥99%) by sonication for 15 min. Extracts were centrifuged at 4500 × g for 5 min, and the supernatant was filtered through a 0.22 µm nylon filter before being diluted by factors of 2 and 20.

Cannabinoid analysis was performed using a Waters Acquity I-Class UPLC system with a photodiode array (PDA) UV detector. Separation was achieved on a Cortecs 1.6 µm, 2.1 mm × 150 mm column (Waters Corporation, Milford, MA, USA) with a Cortecs C18 pre-column (Waters Corporation, 90Å, 1.6 μm, 2.1 mm X 5 mm) maintained at 30°C. The mobile phase consisted of (A) 20 mM ammonium formate (Thermo Fisher Scientific, Optima LC/MS ≥99%), acidified to pH 2.92 with formic acid (MS grade) and diluted with Mili-Q water (Thermo Fisher Scientific) and (B) 100% acetonitrile (Milipore Sigma, OmniSolv® LC/MS ≥99.9%), with a gradient elution program as follows: 0–6.4 min, 76% B; 6.5–8 min, 99% B; 8.1–10 min, re-equilibration at 76% B. The flow rate was set at 0.45 mL/min, and the injection volume was 1 µL. Detection was performed at 228 nm, and quantification was based on 5-point calibration curves (1–100 mg/L) for all cannabinoid standards.

Standard calibration curves were prepared by diluting a mix of cannabinoid standards in 80% methanol. A 500 μL stock solution at 100 mg/L was prepared by combining 50 μL of each individual standard from stock solutions at 1000 mg/L and adjusting the final volume with 80% methanol. Serial dilutions were then performed to generate calibration points at 100, 50, 25, 5, and 1 mg/L. Details of the cannabinoid standards are provided in Supplementary Table 3.

### Quantification of ACC and SAM

A detailed methodological description of standards preparation, derivatization, and chromatographic conditions is provided in the Supplementary Materials and the accompanying parameters can be found in Supplementary Table 4 through Supplementary Table 7. SAM and ACC concentrations were quantified against standard calibration curves prepared from serial dilutions of frozen stock standards, and lysine was included as an internal derivatization control (Supplementary Table 4). Leaf tissues (n = 3 biological replicates per genotype, treatment, and time point) were harvested on day 0 and day 1, flash frozen, and extracted using a trichloroacetic acid (TCA) protocol (Bishop et al., 2024; Sandhu et al., 2024). Approximately 50 mg of leaf tissue was homogenized in 1 mL TCA, centrifuged, and filtered. Supernatants were derivatized using 6-aminoquinolyl-N-hydroxysuccinimidyl carbamate (AQC; Supplementary Table 4) and analyzed via UPLC-MS/MS on a Waters Acquity I-Class UPLC and Xevo TQ-S system (Supplementary Table 5 & Supplementary Table 6). Chromatographic separation was performed on a BEH C18 column, and analytes were detected in positive ESI mode using multiple reaction monitoring (MRM; Supplementary Table 7). SAM and ACC data processing for quantification was performed using TargetLynx V4.1.

### Statistical analysis of ACC and SAM data

Variance stabilizing transformation (VST) of gene expression data was correlated with measured ACC levels using Spearman’s rank correlation within each treatment group. Genes with zero variance were excluded, and outliers (Studentized residuals > ±3.4) were removed. FDR correction was applied to p-values, and only genes with |ρ| ≥ 0.5 and FDR-adjusted p ≤ 0.05 in at least one group were visualized. ACC concentrations were also analyzed using a linear mixed-effects model including treatment, sex, and day as fixed effects, and sample ID nested within genotype as a random effect. Tukey’s HSD post-hoc test was applied to estimated marginal means. SAM was below the limit of quantification in all samples and was not included in downstream analyses. Full statistical methods, R package versions, and model diagnostics are available in the Supplementary Materials.

### ERG Identification

Identification of ERG in *C. sativa* was previously conducted using an *in silico* ortholog analysis by Monthony *et al*. (2024), based on the *C. sativa* cs10 reference genome (RefSeq assembly accession: GCF_900626175.2; Grassa et al. 2021). We improve upon this dataset by applying the same methodology to newly available genomic resources, including the *C. sativa* ‘Pink Pepper’ XX reference genome (NCBI 2023) and the haploid X and Y whole genomes for the genotype ‘Otto II’ (Carey et al. 2024b). Briefly, ERG orthologs were identified using OrthoFinder (Emms and Kelly 2015) and BLASTP (Altschul et al. 1990), as described in Monthony et al. (2024), followed by manual curation. Chromosome maps were generated with the chromoMap (v4.1.1; Anand and Rodriguez Lopez 2022) package in R.

### Whole genome sequencing and variant calling

Fresh leaf tissues obtained from one male (XY) and one female (XY) individual of the three genotypes were used for DNA extraction using the Qiagen DNeasy Plant Mini kit (Qiagen GmbH, Germany). For whole genome sequencing (WGS), six libraries were prepared at the Institut de biologie intégrative et des systèmes (IBIS), Université Laval, QC, Canada. The sequencing was conducted on an Illumina NovaSeq 6000 (Illumina, CA, USA) with 150 bp paired-end reads at the Genome Quebec Service and Expertise Center (CESGQ), Montreal, QC, Canada, with an average coverage of approximately 30x per sample.

Reads were mapped on the *C. sativa* ‘Pink Pepper’ reference genome (NCBI 2023) with *bwa mem* version 7.17 (Li 2013). Variant calling was performed with Fast-WGS (Torkamaneh et al. 2018) and manta version 1.6 (Chen et al. 2016a) for nucleotide and structural variant calling, respectively. Raw variant data were filtered with VCFtools version 1.16 (Danecek et al. 2011) to remove low-quality variants (filter PASS, QUAL <10, MQ <30, MinDP <6, MAF 1%). Variants within ±5 kb of CsERGs in the *C. sativa* ‘Pink Pepper’ reference genome (NCBI 2023) were filtered using BEDtools v2.31.1 (Quinlan and Hall 2010). Functional impact of variants in ±5 kb around of CsERGs were investigated using the Ensembl Variant Effect Predictor (McLaren et al. 2016) with the GFF of Pink Pepper (NCBI 2023). Consequence terms (e.g., inframe insertion, missense variant, synonymous variant) and the impact rating are based on Sequence Ontology (Eilbeck et al. 2005). Impact ratings are classified as (i) Modifier correspond to variant affecting non-coding sequences, (ii) Low correspond to variant unlikely to change protein behaviour, (iii) Moderate correspond non-disruptive variant that might change protein effectiveness, and (iv) High correspond to variants probably causing protein truncation and/or loss of function. The nucleotide diversity (π; Nei & Li, 1979) in a sliding windows of 1000 bp and Tajima’s D value in bins with size of 10,000 SNPs were estimated using VCFtools (Danecek et al. 2011). Gene with less than 2 SNPs were excluded from the diversity analysis.

### RNA extraction & sequencing

Leaf tissues were consistently sampled from mature leaves, in the middle section of the plant. Representative floral tissues (day 14) were consistently sampled from upper most axillary branch of the plants. Immediately after harvesting, plant tissues were flash-frozen in liquid nitrogen and hand-ground using a disposable cryosafe pestle in 2 mL RNase-free tubes to preserve RNA integrity. Total RNA was extracted using the RNeasy Plant Mini Kit (Qiagen), following the manufacturer’s protocol with modifications to enhance RNA purity. Briefly, 25–50 mg of ground, frozen tissue was combined with RLC buffer and β-mercaptoethanol, as per the manufacturer’s recommendations, and vortexed until homogenized. To improve purity, an additional wash with buffer RPE (Qiagen) was included in the final wash step. RNA was quantified spectrophotometrically using a NanoDrop spectrophotometer (Thermo Fisher Scientific). RNA quality was assessed visually, via electropherograms from the 2100 Bioanalyzer (Agilent Technologies, Santa Clara, CA, USA), and quantitatively, using the RNA Integrity Number (RIN; Supplementary Table 8). High-quality RNA samples were converted to cDNA using the NEBNext® Ultra™ II Directional RNA Library Prep Kit for Illumina (New England Biolabs, Ipswich, MA, USA). Library preparation and sequencing were conducted at the Institut de Biologie Intégrative et des Systèmes (IBIS), Université Laval, Quebec, Canada. Sequencing was performed on an Element Biosciences AVITI platform (Element Biosciences, San Diego, CA, USA), generating 150 bp paired-end reads, with an average output of ∼30 million reads per sample. In total, 138 samples passed quality control and were used in all future analyses (Supplementary Table 8).

### Transcriptomic analysis

RNA sequencing quality was assessed using FASTQC (v0.11.8), and trimming of low-quality bases was performed in paired-end mode using Trimmomatic (v0.39) with the following parameters: adapter trimming (ILLUMINACLIP), a mismatch seed value of 2, a palindrome clip threshold of 30, a simple clip threshold of 10, and a minimum read length (MINLEN) of 40. For rare RNA samples identified with foreign RNA contamination based on overrepresented sequences and skewed GC ratios, contaminant sequences were extracted for further analysis. These sequences were classified by species using a BLAST search against the NCBI nucleotide database, with duplicates filtered and mapped to unique organisms. Non-plant contaminants, such as animal and fungal sequences, were removed using a modified Trimmomatic script with a custom contaminant sequence list. The trimming process retained ∼95% of paired reads, and post-trimming quality was re-assessed using FastQC to confirm improvements in sequence composition.

Alignment of paired-end reads was performed as described in Monthony et al. (2024) with modifications. Briefly, reads were aligned to the *C. sativa* “Pink Pepper” v1 genome (NCBI) using STAR (v2.7.11b) and output as coordinate-sorted BAMs (--outSAMtype BAM SortedByCoordinate). Immediately after each sample finished mapping, they were aligned with Samtools (v1.15) to retain only alignments with MAPQ ≥ 30 (samtools view -b -q 30), yielding high-confidence BAMs for downstream counting. Gene-level counts were then obtained with HTSeq-count (v2.0.2; Putri et al. 2022) in positional mode (-r pos), using a stranded setting (-s yes), a minimum alignment quality filter of 30 (-a 30), and exon features (-t exon) keyed by gene IDs (-i gene).

### Principal component analysis and gene expression visualization

Principal component analysis (PCA) was performed on the VST normalized, batch-corrected counts of 47 selected genes of interest, obtained from gene-level counts following alignment and quantification. VST transformation was applied using the DESeq2 package (v1.42.1; Love et al. 2014) to normalize the gene expression data. The first two principal components (PC1 and PC2) were extracted and plotted to visualize the clustering of samples based on their gene expression profiles. PCA plots were generated using the ggplot2 package (v3.5.1), with ellipses representing 95% confidence intervals for each treatment group.

For heatmap preparation, the filtered gene expression data was scaled by row (z-score transformation) using the pheatmap package (v1.0.12) to normalize gene expression levels across samples. Hierarchical clustering of genes and samples was performed using Euclidean distance and complete linkage to reveal patterns in gene expression across different treatments and time points.

### Network analysis (WGCNA)

Weighted Gene Co-expression Network Analysis (WGCNA) was performed to construct sex-specific gene co-expression networks, identifying gene expression patterns associated with sex-change treatments. All analyses were performed in R (v4.3.3) using the WGCNA package (v1.73; Langfelder and Horvath 2008). Networks were built separately for male (XY) and female (XX) samples, using count data processed through multiple quality control and normalization steps to ensure data integrity.

Prior to network construction, outlier genes and samples were detected and removed using the goodSamplesGenes (gsg) function in WGCNA, which identifies genes and samples with excessive missing values or abnormal expression patterns (see Supplementary Table 8). To further validate sample quality, hierarchical clustering and principal component analysis (PCA) were used to identify additional outlier samples, which were subsequently removed. Following outlier removal, batch correction was performed using a DESeq2-based method (Roy et al. 2025). Additionally, low-count genes were filtered using HTSFilter (v1.42.0; Rau et al. 2013), which dynamically adjusts filtering thresholds based on dataset structure rather than applying an arbitrary cutoff. After filtering, the data were normalized using variance stabilizing transformation (VST) to reduce heteroscedasticity and improve network construction.

To determine an appropriate soft-thresholding power, the pickSoftThreshold function was applied independently to each subset to ensure scale-free topology (signed R² > 0.80 with minimal mean connectivity). Based on topology evaluation, a soft-thresholding power of 6 was selected for XX and 7 for XY datasets. Networks were constructed using blockwiseModules, employing a signed adjacency matrix, a module merging cut height of 0.20, with a maximum block size of 30,000 and modules were identified using dynamic tree cutting.

Module-trait correlation, adapted from Wang *et al*. (2020), was performed independently for XX and XY networks to identify biologically relevant gene clusters associated with sex-altering treatment. Treatment was defined as STS application in the XX network and ethephon application in the XY network. To prioritize biologically relevant gene modules, modules significantly correlated with treatment (|r| ≥ 0.5, p ≤ 0.05) were retained for further analysis (Wang et al. 2020). To assess whether treatment-associated modules exhibited temporal specificity, their correlation with time points was examined. Time points were analyzed using a “Day vs. all” approach, where Day 1 and Day 14 were each compared to all other time points. Day was treated as a categorical variable with Day 0 as the baseline, while treatment was encoded as a binary variable (1 = treated, 0 = control). A shortlist of ethylene-related genes (ERGs) was generated by extracting those present in highly correlated modules. For each ERG, intra-modular connectivity (kME) was calculated to assess its centrality within its respective module. The final list of ERGs from highly correlated modules in both XX and XY networks was retained for subsequent differential gene expression analysis.

### Differential gene expression analysis

Differential gene expression (DGE) analysis was performed using DESeq2 to identify genes exhibiting significant expression changes in response to ethephon and STS treatments across male (XY) and female (XX) plants, respectively. DGE was focused on candidate ERGs identified in WGCNA. Raw count data were preprocessed by filtering low-expression genes using HTSFilter (Rau et al. 2013). Normalized counts were obtained using a VST. Batch effects were modeled as a fixed effect in the DESeq2 generalized linear model (GLM) to correct for potential variation across experimental batches. Pairwise contrasts were constructed to evaluate: (1) STS-treated IMF (XX) vs. untreated FF (XX), (2) ethephon-treated IFF (XY) vs. untreated MF (XY), and (3) baseline differences between control MF (XY) and FF (XX) samples. Custom contrast vectors were applied to account for sex, treatment, and batch effects, ensuring robust comparisons. The log_2_ fold-change (log_2_FC) for each contrast was shrunk using ashr (Stephens 2017) to provide more stable effect size estimates, particularly for low-expression genes.

Genes were considered significantly differentially expressed if they met the criteria of adjusted p-value (padj) ≤ 0.05 and absolute log_2_FC ≥ 1 (Monthony et al. 2024; Shi et al. 2024b). Differentially expressed genes were further filtered to only include those differentially expressed across all three genotypes, ensuring treatment effects were not genotype dependent. Visualization of DGE results included bar plots of log_2_FC values, highlighting genes with significant changes across treatment conditions and time points. Additionally, variance-stabilized expression levels were plotted using boxplots to compare expression profiles across MF, FF, IMF, and IFF groups. All analyses were conducted using R (v4.3.3), and visualizations were produced using ggplot2 (v3.5.1) for data visualization, dplyr (v1.1.4) for data manipulation, and wesanderson (v0.3.7) for color palettes.

## Statements and Declarations

The authors have no competing interests to declare. The authors gratefully acknowledge the support of the Natural Sciences and Engineering Research Council (NSERC) Discover Grant number RGPIN-2022-03396. ASM has also been supported by a NSERC Canada Vanier Graduate Scholarship.

## Data availability statement

All raw transcriptomic data will be made available on the NCBI sequence read archive (SRA) upon publication of this manuscript.

## Acknowledgements

The authors would like to thank Christian Otis and Brian Boyle at the Université Laval Genomic Analysis Platform (PAG) for their invaluable counsel on RNA extraction and Hélène Martin for her help sexing plant materials used in this study.

## Author contributions

ASM and DT planned and designed the research. ASM, MdR, OC and SJM contributed to the methodology. ASM, JR, MdR and OC performed the investigation and contributed to data curation. ASM, JR, MdR conducted formal analysis. SJM and DT acquired funding for the project. ASM and DT handled project administration. SJM and DT provided resources. SJM and DT supervised the research. ASM, MdR and JR visualized the results. ASM, MdR, OC and DT wrote the original draft of the manuscript. ASM, SJM, and DT contributed to reviewing and editing the manuscript.

## Supplementary Table Captions

**Supplementary Table 1** Cutting, vegetative and flowering growth conditions. Following rooting of cuttings, all plants were grown in controlled growth chambers. Male flowers (MF); female flowers (FF); induced female flowers (IFF); induced male flowers (IMF); as needed (PRN), Photosynthetic Photon Flux Density (PPFD).

**Supplementary Table 2** Primer sequences used for sexing in a PACE assay, table extracted from Toth (2022).

**Supplementary Table 3** Cannabinoid standards used in the cannabinoid quantification by UPLC-PDA. All standards were purchased from Agilent.

**Supplementary Table 4** ACC and SAM working solutions (WS) and calibration standards (C) concentrations before/after derivatization with AccQ-Tag.

**Supplementary Table 5** UPLC Gradient Elution Program Adapted from Kairos et al. (2019). The table outlines the gradient conditions used for chromatographic separation of ACC and SAM. Mobile phase A consists of 0.1% formic acid in water, while mobile phase B consists of 0.1% formic acid in acetonitrile.

**Supplementary Table 6** Optimized Xevo TQ-S Mass Spectrometry Parameters for ACC and SAM Analysis.

**Supplementary Table 7** Multiple Reaction Monitoring (MRM) Transitions for ACC and SAM Detection. m/z: refers to the mass-to-charge ratio of the ions detected; sd: denotes single dissociation, indicating that the precursor ion undergoes a single-step fragmentation before detection.

**Supplementary Table 8** Metadata and RNA-seq quality control metrics for samples included in this study. Columns include sample name, genotype, genetic sex, tissue type, sample timepoint, treatment (Ctrl = control; ET = ethephon; STS = Silver thiosulfate), RNA integrity number (RIN) following sample RNA extraction, sequencing depth (coverage, ×), GC content (%), and average mapping quality following alignment with STAR and post-alignment QC using Qualimap. The final two columns indicate whether each sample passed internal quality filters and was retained for downstream weighted gene co-expression network analysis (WGCNA) and differential expression analysis (DESeq2), respectively.

**Supplementary Table 9** Results of the F-test from the ANOVA for the percentage of *C. sativa* florets which underwent sex change.

**Supplementary Table 10** Cannabinoid content (% w/w) of mature female flowers for the genotypes used in the present study and the analytical limit of quantification (LOQ) and limit of detection (LOD) for the cannbinoids analyzed. ND indicates that the sample was below the limit of detection. The cannabinoid abbreviations are as follows: tetrahydrocannabinolic acid (THCA), delta-9-tetrahydrocannabinol (Δ9-THC), cannabidiolic acid (CBDA), cannabidiol (CBD), cannabigerolic acid (CBGA), cannabigerol (CBG), cannabichromene (CBC), cannabinol (CBN), tetrahydrocannabivarin (THCV), and cannabidivarin (CBDV).

**Supplementary Table 11** *C. sativa* ERGs as identified through homology analysis using the current reference genome (‘Pink Pepper’; NCBI, 2023), X and Y haplotypes from the Otto II genotype, and the previous reference genome, Cs10 (Grassa et al. 2021). Chr: chromosome number.

**Supplementary Table 12** Genetic diversity analysis of XX and XY genotypes Deadly Kernel, La Rosca and Panama Pupil V4.

**Supplementary Table 13** Quantification data (ug/g fresh weight) of 1-aminocyclopropane-1-carboxylic acid (ACC) and detection of S-adenosylmethionine (SAM) by UPLC-MS.

**Supplementary Table 14** ERGs identified within treatment-associated modules from the XX and XY network before differential gene expression filtering. Module-trait correlations (r) indicate the relationship between each module and treatment (STS for XX and ethephon for XY) or time points (Day 1 and Day 14). Intramodular connectivity (kME) reflects the centrality of each gene within its assigned module.

## Supplementary Figures Captions

**Supplemental Figure 1** Representative timeline of floral development in MF, IMF, FF and IFF of *C. sativa* ‘Deadly Kernel’ (DK) and ‘Panama Pupil V4’ (PP) from 7 to 28 days after the photoperiod shift from long day to short day. N/A indicates photo not available for a given time/treatment.

**Supplemental Figure 2** Differential expression data contrasting control (MF or FF) and sex changed (IFF or IMF) plants at day 1 and 14 using the 24 ERGs identified from WGCNA. Genes which show genotype-independent behaviour (gray highlight) were retained for further analysis.

**Supplemental Figure 3** Genomic distribution and predicted impact of SNPs and structural variants (SVs) in the 47 ERGs of the six C. sativa samples. A) Schematic representation of a gene structure and its surrounding regulatory regions (5 kb upstream and downstream), highlighting genomic elements considered in the analysis. Regions are color-coded based on their predicted impact, ranging from modifier (low impact, mostly regulatory) to low, moderate, or high impact (potential functional consequences protein function). Distribution of SNPs (B) and SVs (C) categorized by their predicted impact and localization. D) Distribution of modifier impact variants (SNPs and SVs) across the 47 ERGs. E) Distribution of low to high impact variants identified in the 47 ERGs.

**Supplemental Figure 4** Endogenous ACC concentrations and their correlation with CsCTR1-LIKE expression in C. sativa leaves during floral transition. A) ACC quantification at vegetative (day 0) and flowering (day 1) stages across experimental groups. Error bars represent standard error of the estimated marginal means. Different letters indicate significant differences among groups (Tukey’s HSD, α = 0.05). B) Spearman correlation heatmap depicting the relationship between CsCTR1-LIKE (LOC115706236) expression and ACC concentration across treatment groups. A significant negative correlation (ρ = −0.8, p ≤ 0.001) is observed in XX Day 0 – FF. Statistical significance for Spearman correlations was categorized as follows: p ≤ 0.001, ***; p ≤ 0.01, **; p ≤ 0.05, *; correlations with p > 0.05 were considered non-significant.

**Supplemental Figure 5** WGCNA module identification and correlation analysis. A, B) Clustering dendrogram for XX (A) and XY (B) plants. C-D) Module-trait correlation heatmaps of identified modules correlated with treatment and day. Modules significantly associated with the traits with identified with |cor | > 0.5 and p value ≤ 0.05, and are indicated by asterisks as follows: p ≤ 0.001, ***; p ≤ 0.01, **; p ≤ 0.05, *; correlations with p > 0.05 were considered non-significant. Red and blue color notes positive and negative correlation between a given module and trait pair.

## Notes

### Competing Interest Statement

The authors have declared no competing interest.

### Summary of Updates

Writing revisions to improve readability, clarify and brevity were done in the following sections: Title Abstract Discussion Supplementary tables were combined to reduce the overall number of tables.

