## Supplementary Materials for "Sex-Specific Ethylene Responses Drive Floral Sexual Plasticity in Cannabis"

### Supplementary Figures

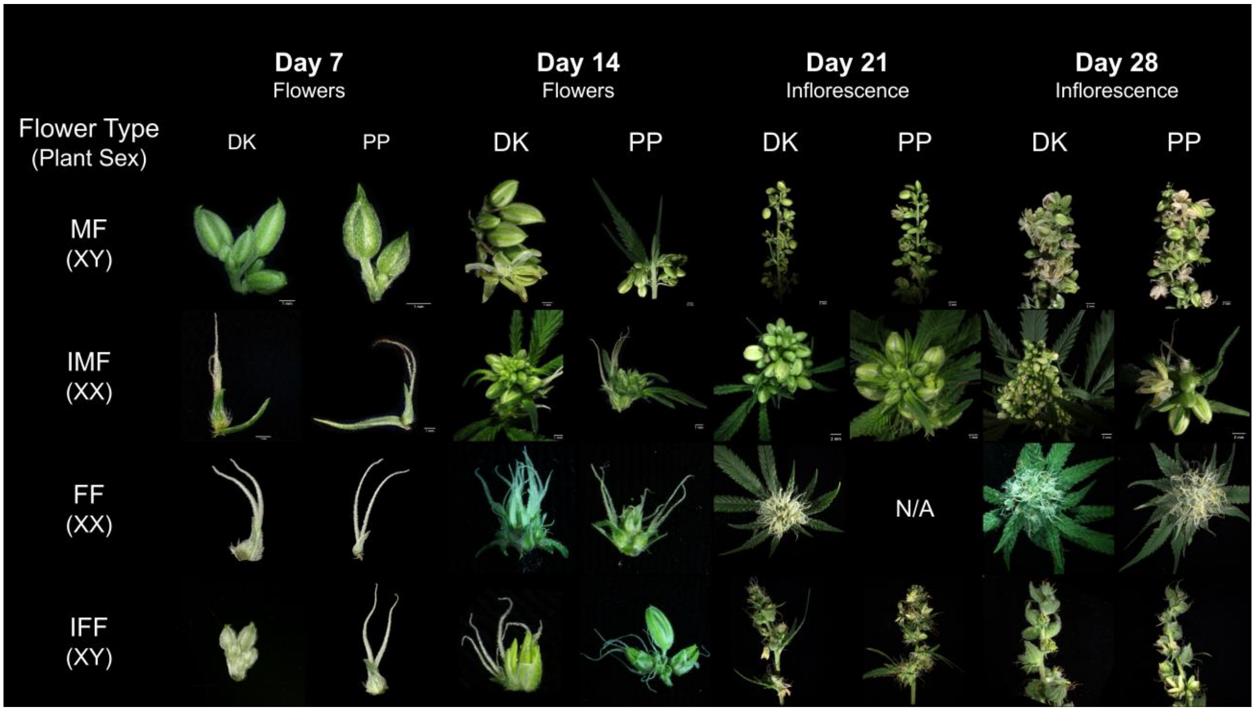


**Supplemental Figure 1** Representative timeline of floral development in MF, IMF, FF and IFF of *C. sativa* 'Deadly Kernel' (DK) and 'Panama Pupil V4' (PP) from 7 to 28 days after the photoperiod shift from long day to short day. N/A indicates photo not available for a given time/treatment.

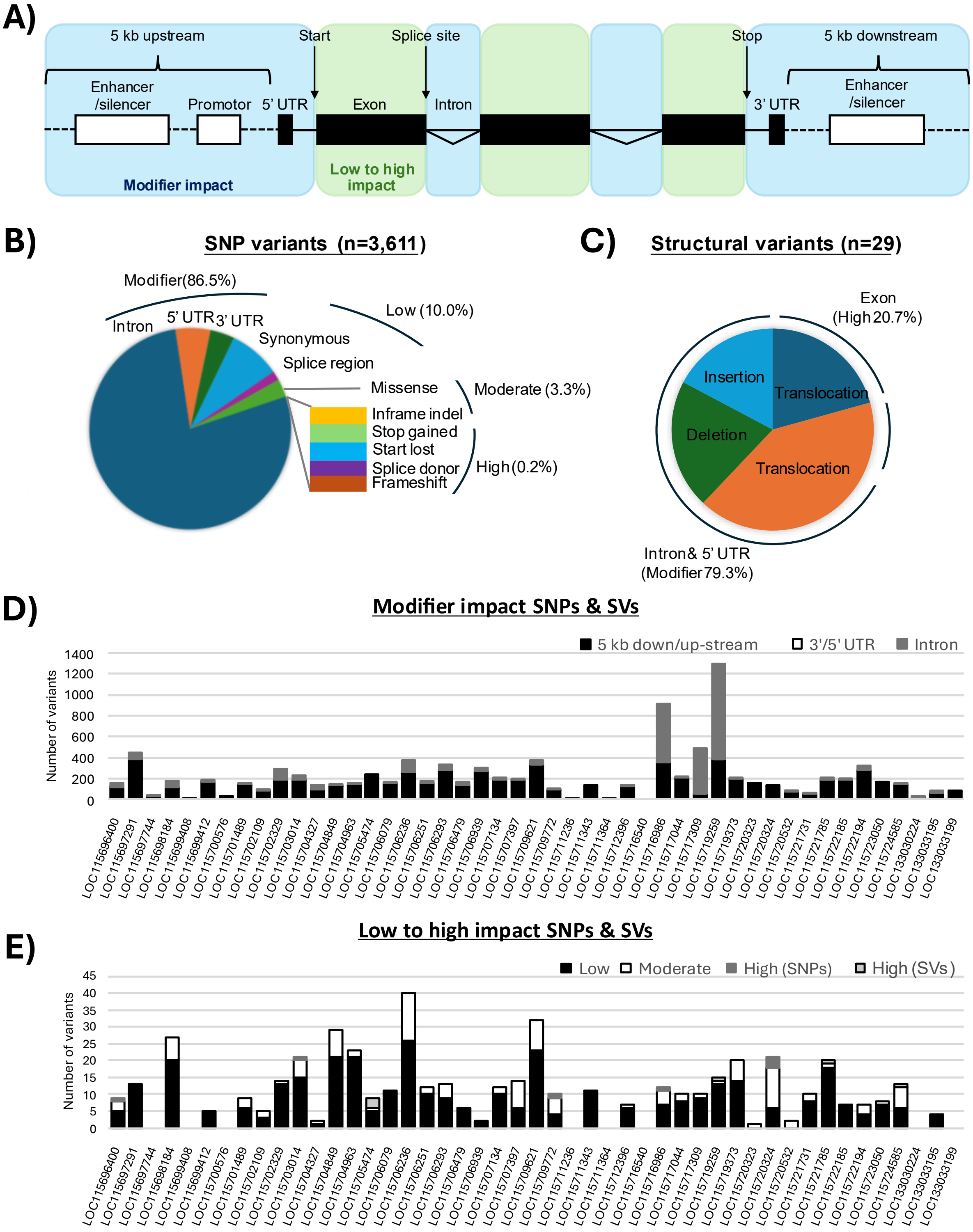
A)
5 kb upstream
Start
Splice site
Stop
5 kb downstream
Enhancer /silencer
Promotor
5' UTR
Exon
Intron
3' UTR
Enhancer /silencer
Modifier impact
Low to high impact
B)
SNP variants (n=3,611)
Modifier(86.5%)
Intron
5' UTR
3' UTR
Synonymous
Splice region
Low (10.0%)
Missense
Moderate (3.3%)
Inframe indel
Stop gained
Start lost
Splice donor
Frameshift
High (0.2%)
C)
Structural variants (n=29)
Exon (High 20.7%)
Insertion
Translocation
Deletion
Translocation
Intron& 5' UTR (Modifier 79.3%)
D)
Modifier impact SNPs & SVs
5 kb down/up-stream
3'/5' UTR
Intron
Number of variants
E)
Low to high impact SNPs & SVs
Low
Moderate
High (SNPs)
High (SVs)
Number of variants

**Supplemental Figure 2** Genomic distribution and predicted impact of SNPs and structural variants (SVs) in the 47 ERGs of the six C. sativa samples. A) Schematic representation of a gene structure and its surrounding regulatory regions (5 kb upstream and downstream), highlighting genomic elements considered in the analysis. Regions are color-coded based on their predicted impact, ranging from modifier (low impact, mostly regulatory) to low, moderate, or high impact (potential functional consequences protein function). Distribution of SNPs (B) and SVs (C) categorized by their predicted impact and localization. D) Distribution of modifier impact variants (SNPs and SVs) across the 47 ERGs. E) Distribution of low to high impact variants identified in the 47 ERGs.

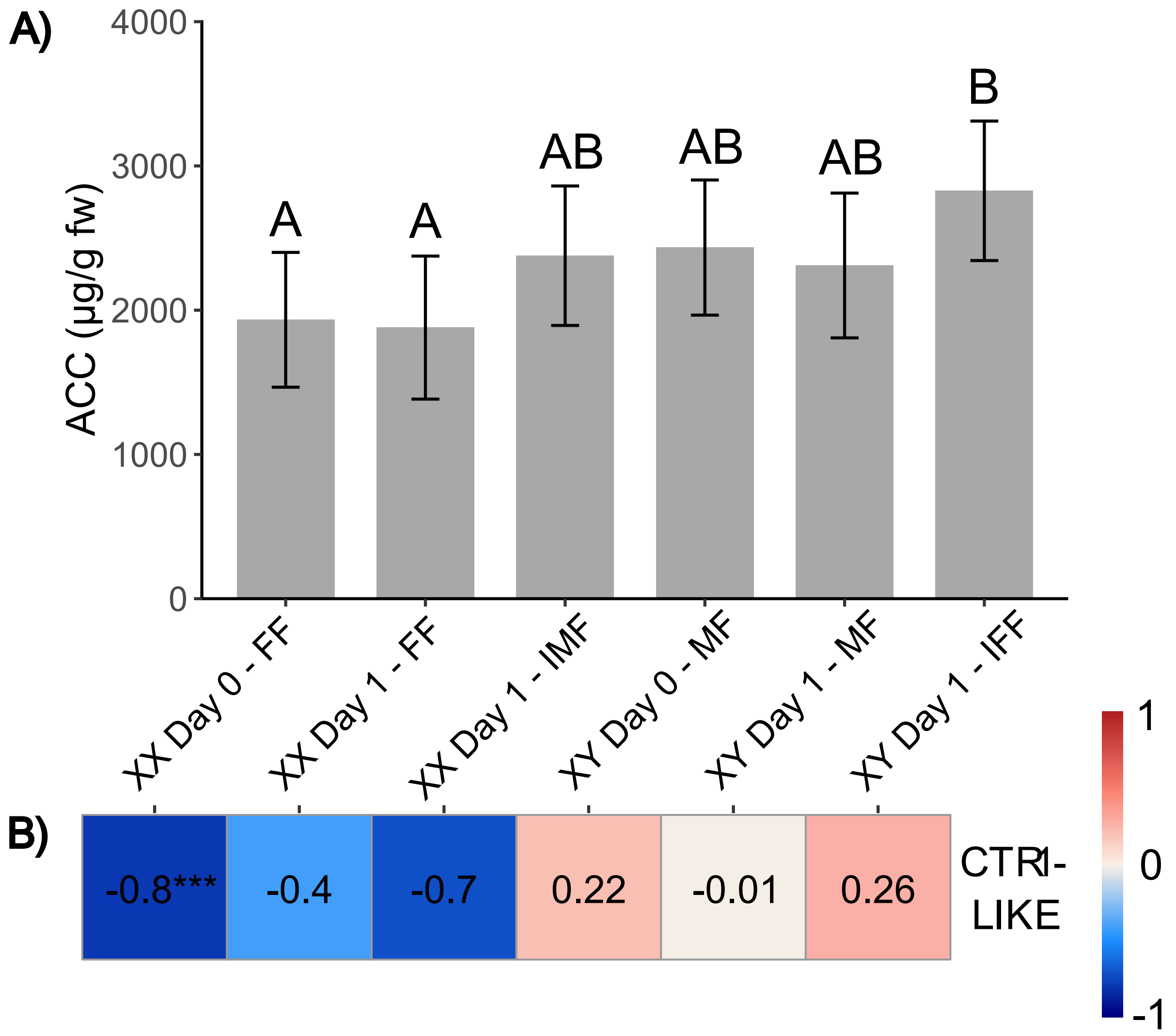

**Supplemental Figure 3** Endogenous ACC concentrations and their correlation with CsCTR1-LIKE expression in C. sativa leaves during floral transition. A) ACC quantification at vegetative (day 0) and flowering (day 1) stages across experimental groups. Error bars represent standard error of the estimated marginal means. Different letters indicate significant differences among groups (Tukey's HSD, $\alpha = 0.05$). B) Spearman correlation heatmap depicting the relationship between CsCTR1-LIKE (LOC115706236) expression and ACC concentration across treatment groups. A significant negative correlation ($\rho = -0.8$, $p \leq 0.001$) is observed in XX Day 0 – FF. Statistical significance for Spearman correlations was categorized as follows: $p \leq 0.001$, ***; $p \leq 0.01$, **; $p \leq 0.05$, *; correlations with $p > 0.05$ were considered non-significant.

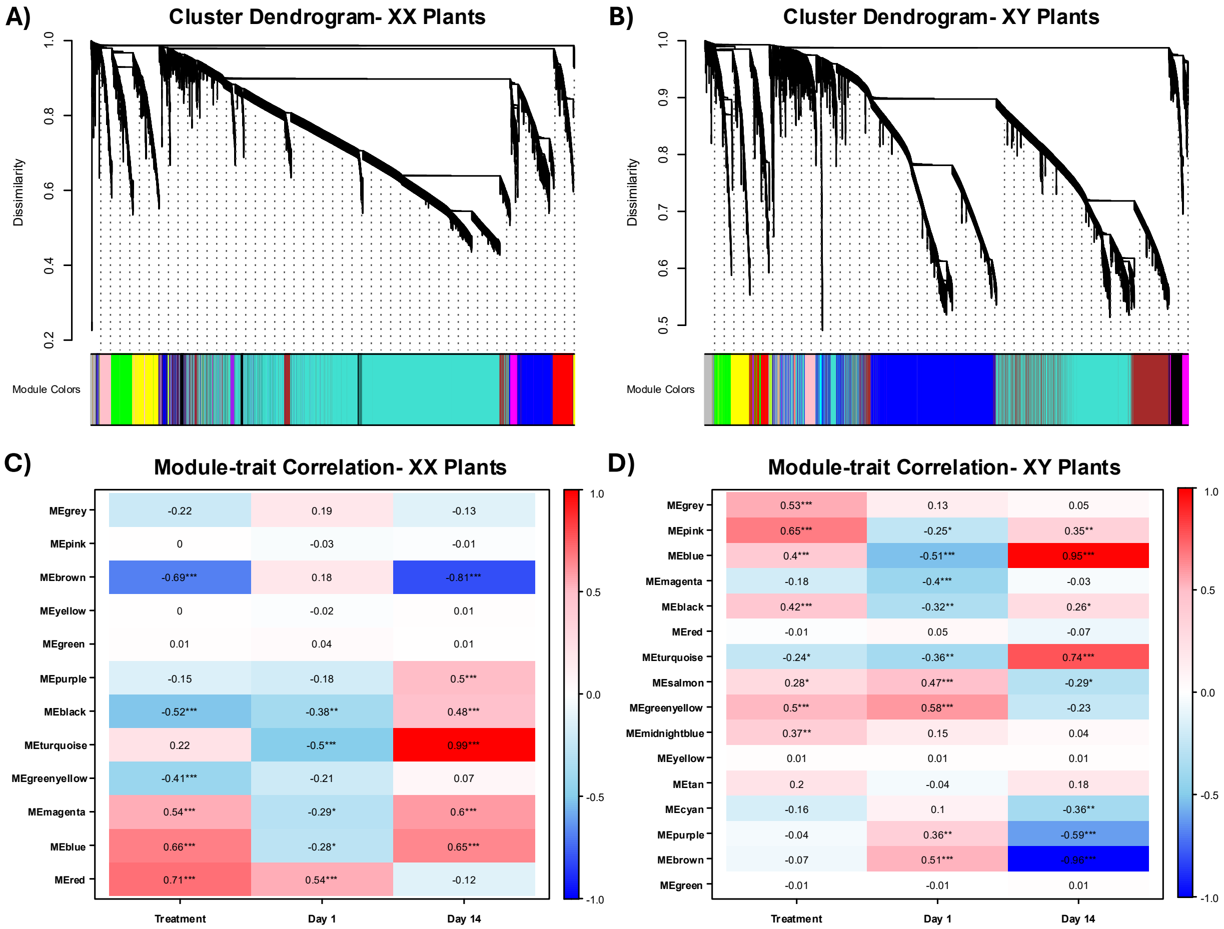


**Supplemental Figure 4** WGCNA module identification and correlation analysis. A, B) Clustering dendrogram for XX (A) and XY (B) plants. C-D) Module-trait correlation heatmaps of identified modules correlated with treatment and day. Modules significantly associated with the traits with identified with $|cor| > 0.5$ and p value $\leq 0.05$, and are indicated by asterisks as follows: $p \leq 0.001$, ***; $p \leq 0.01$, **; $p \leq 0.05$, *; correlations with $p > 0.05$ were considered non-significant. Red and blue color notes positive and negative correlation between a given module and trait pair.

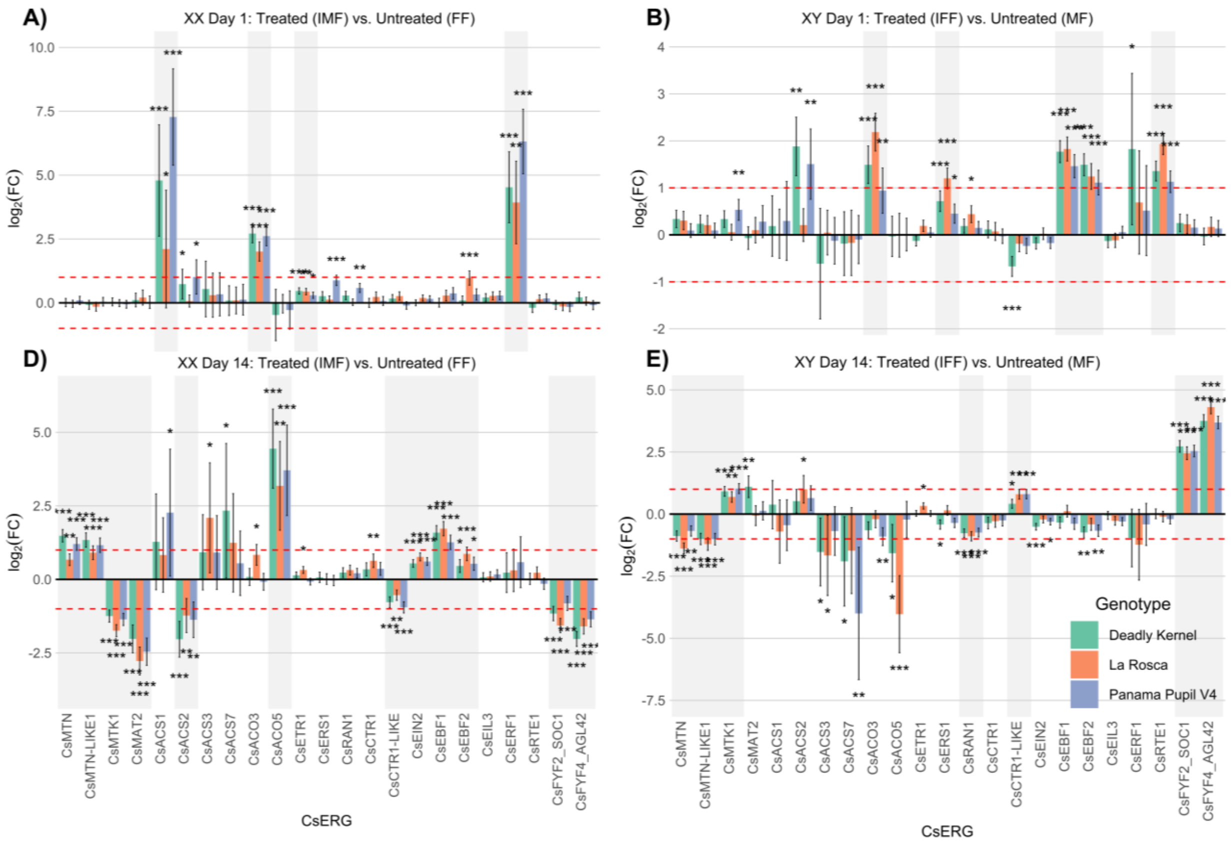


**Supplemental Figure 5** Differential expression data contrasting control (MF or FF) and sex changed (IFF or IMF) plants at day 1 and 14 using the 24 ERGs identified from WGCNA. Genes which show genotype-independent behaviour (gray highlight) were retained for further analysis.

### Supplementary Information

##### Preparation of ACC and SAM Standards

Stock solutions of 1-aminocyclopropanecarboxylic acid (ACC; Sigma) and S-(5′-Adenosyl)-L-methionine chloride dihydrochloride (SAM; Sigma) were prepared using ultrapure water and 0.1 N HCl. For ACC, 0.2528 g was dissolved in 25 mL of ultrapure water (filtered through a 0.45 µm Nylon filter) to obtain a 0.1 M stock solution. A 2.5 mM working stock solution was then prepared by adding 625 µL of the 0.1 M ACC stock to 25 mL of 0.1 N HCl, followed by serial dilutions to obtain progressively lower concentrations (Supplementary Table 4). For SAM, 100 mg was dissolved in 1 mL of 0.1 N HCl, transferred to a 2.0 mL Eppendorf Safe-Lock tube, and rinsed with an additional 1 mL of 0.1 N HCl, yielding a final stock concentration of 50 mg/mL (0.1255 M). Subsequent dilutions were performed to generate 5.09 mg/mL and 0.509 mg/mL solutions. To prevent degradation due to repeated freeze-thaw cycles, SAM standards were stored at -80°C in aliquots. A 10,000 µM (3.98 mg/mL) working standard solution was prepared from a thawed 50 mg/mL aliquot and stored at -80°C in separate 100 µL aliquots. All ACC working solutions were stored in Corning microcentrifuge tubes with sealable screw caps (VWR) at -20°C.

##### ACC and SAM Calibration curve preparation

A calibration curve was prepared daily using frozen ACC and SAM aliquots. The 10,000 µM SAM standard was retrieved from the -80°C freezer, while the 250 µM ACC standard was taken from -20°C. After thawing at room temperature for approximately 10 minutes, 50 µL of SAM, 10 µL of ACC, and 40 µL of 0.1 N HCl were combined to form a 100 µL calibration solution (C1). A series of serial dilutions was then prepared by transferring 75 µL of C1 into 75 µL of 0.1 N HCl to create the C2 standard. This process was repeated sequentially to generate a total of six calibration standards (Supplementary Table 5).

#### Sample preparation and derivatization

Sample preparation followed an established trichloroacetic acid (TCA) extraction protocol (Bishop *et al.*, 2024; Sandhu *et al.*, 2024). Leaf tissue samples were collected on day 0 and day 1, with three biological replicates per genotype, treatment, and time point and flash frozen using liquid nitrogen, until analysis. Approximately 50 mg of each sample, avoiding major veins, was homogenized in 1 mL of TCA using a blue stick homogenizer for 5 minutes, with intermittent vortexing to ensure complete tissue disruption. The homogenized mixture was then centrifuged at 13,000 × g for 5 minutes, and approximately 800 µL of the resulting supernatant was transferred to a centrifuge filter tube. To further clarify the extract, a second centrifugation was performed at 13,000 × g for 5 minutes, completing the filtration process.

Extracted samples and standards were derivatized with 6-aminoquinolyl-N-hydroxysuccinimidyl carbamate (AQC) derivatization reagent (Waters AccQ Tag Ultra reagent kit; Waters Inc., Mississauga, ON). For each derivatization reaction, 70 µL of borate buffer was added to a conical glass insert within an amber HPLC vial, followed by 10 µL of the sample or standard solution and 20 µL of AccQ-Tag reagent (Supplementary Table 5). The mixture was vortexed for 5 seconds, left to stand at room temperature for 1 minute, and heated at 55°C for 10 minutes to complete the derivatization process.

**UPLC-MS analysis**

The UPLC-MS method used in this study was adapted from Sandhu et al. (2024). Derivatized samples were analyzed using a Waters Acquity I-Class UPLC system coupled to a Waters Xevo TQ-S Triple Quadrupole Mass Spectrometer (Waters Inc., Mississauga, ON). Chromatographic separation was performed using an Acquity reverse-phase column (150 × 2.1 mm, 1.7 µm C18 BEH), with a 10 µL injection volume and a flow rate of 0.4 mL/min. The mobile phase consisted of 0.1% formic acid (FA) in water (mobile phase A) and 0.1% FA in acetonitrile (mobile phase B), with elution performed according to a gradient method detailed in Supplementary Table 6. To prevent carryover, the system was equipped with an automated needle washing routine, using 0.1% FA in water (weak wash) and 0.1% FA in acetonitrile (strong wash), while a seal wash of 80:20 water/methanol was employed to maintain system integrity.

Tandem mass spectrometry was conducted in positive electrospray ionization (ESI+) mode, with optimized instrument parameters provided in Supplementary Table 7. Multiple reaction monitoring (MRM) transitions were used to quantify ACC and SAM, with lysine included as an internal control to verify derivatization efficiency (Supplementary Table 8). Data acquisition and quantitative analysis were performed using TargetLynx V4.1 software (Waters Inc., Mississauga, ON). The software was configured to integrate MRM transitions, with retention times predicted based on calibration standards and peak integration performed within a defined retention time window (Supplementary Table 9).

**Data analysis**

To assess the relationship between ethylene-related gene expression and ACC accumulation, Spearman's rank correlation coefficient (Auslander *et al.*, 2016; Qin *et al.*, 2022) was computed between variance-stabilized transformed (VST) gene expression levels and ACC concentrations within each sex, day, and treatment group. Prior to analysis, Studentized residuals exceeding ±3.4 were identified as extreme outliers and removed to prevent undue influence on correlation estimates. Genes exhibiting zero variance within any group were excluded from the analysis. Correlation coefficients (ρ) and associated p-values were computed for each gene within each group, and Benjamini-Hochberg false discovery rate (FDR) correction was applied to adjust for multiple comparisons. Genes meeting the significance threshold ($|\rho| \geq 0.5$, FDR-adjusted $p \leq 0.05$) in at least one group were retained across all groups for visualization. Gene-ACC correlations were visualized across groups using a heatmap generated with *pheatmap* (v1.0.12; Kolde, 2018) in R and significant correlations were annotated with asterisks.

Differences in ACC concentrations between experimental groups were analyzed using a linear mixed-effects model (LME) package *nlme* (v3.1-166; Pinheiro *et al.*, 2025) in R according to the methods described by Monthony *et al.* (Monthony *et al.*, 2025). The model included treatment, sex, and day as fixed effects, with sample ID nested within genotype as a random effect to account for genetic variability. Model residuals were assessed for normality (Shapiro-Wilk test) and variance homogeneity (Levene's test). Tukey's HSD post-hoc test was conducted on estimated marginal means (*emmeans* v1.10.5; Lenth, 2017). Groups with statistically distinct ACC concentrations were annotated using compact letter displays (CLD) to indicate significance groupings (α = 0.05). Bar plots were generated to display mean ACC concentrations (µg/g FW) with standard error of the mean (SEM) for each group.

#### Supplementary Table Captions

*Supplementary Tables can be found in the Excel file accompanying this manuscript.*

**Supplementary Table 1** Cutting, vegetative and flowering growth conditions. Following rooting of cuttings, all plants were grown in controlled growth chambers. Male flowers (MF); female flowers (FF); induced female flowers (IFF); induced male flowers (IMF); as needed (PRN), Photosynthetic Photon Flux Density (PPFD).

**Supplementary Table 2** Primer sequences used for sexing in a PACE assay, table extracted from Toth (2022).

**Supplementary Table 3** Cannabinoid standards used in the cannabinoid quantification by UPLC-PDA. All standards were purchased from Agilent.

**Supplementary Table 4** ACC and SAM standard concentrations. WS: Working solution.

**Supplementary Table 5** ACC and SAM Calibration Range Before/After Derivatization with AccQ-Tag.

**Supplementary Table 6** UPLC Gradient Elution Program Adapted from Kairos et al. (2019). The table outlines the gradient conditions used for chromatographic separation of ACC and SAM. Mobile phase A consists of 0.1% formic acid in water, while mobile phase B consists of 0.1% formic acid in acetonitrile.

**Supplementary Table 14** Genetic diversity analysis of XX and XY genotypes Deadly Kernel, La Rosca and Panama Pupil V4.

**Supplementary Table 15** Quantification data (ug/g fresh weight) of 1-aminocyclopropane-1-carboxylic acid (ACC) and detection of S-adenosylmethionine (SAM) by UPLC-MS.

**Supplementary Table 16** ERGs identified within treatment-associated modules from the XX and XY network before differential gene expression filtering. Module-trait correlations (r) indicate the relationship between each module and treatment (STS for XX and ethephon for XY) or time points (Day 1 and Day 14). Intramodular connectivity (kME) reflects the centrality of each gene within its assigned module.
